# AI-HEAD: Auto-indexing for Helical assembles’ diffraction and fast dynamic structure determination

**DOI:** 10.1101/2025.11.17.688965

**Authors:** Taining Pei, Mingyang Yu, Song Li, Danyang Zhang, Fei Sun, Yan Zhang

**Author notes:** Correspondence should be addressed to Danyang Zhang, Fei Sun, Yan Zhang.

## Abstract

Analyzing the structure of the helical assembly is very important to excavate the mechanism of base life activities such as cell morphology maintaining, cargo transportation, et. al. However, the structure determination of helical assembles still challenging due to ambiguous of helical parameters. Here, we developed automatic, fast and accurate helical parameter and structure calculation method AI-HEAD, which perform diffraction indexing and parameter calculation automatically in seconds, and determine the final structure unambiguously by sorting parameters semi-automatically. AI-HEAD is open source and also provides a user-friendly graphical interface.

## Main

Single particle analysis (SPA) technology of cryo-electron microscopy (cryo-EM) utilizes multiple two-dimensional observations of a protein in different orientations for three-dimensional reconstruction. This method is equally applicable to the structural determination of helical assemblies. As early as 2007, Edward successfully developed the Iterative Helical Real-Space Reconstruction (IHRSR) method (Egelman, 2007) for resolving helical structures using SPA, which has since been widely adopted. Today, the number of helical structures deposited in the EMDB (Electron Microscopy Data Bank) continues to grow annually, with progressively higher resolutions being achieved. The 3D helical reconstruction is based on the assumption that each subunit within a assemble has a defined orientation and position with respect to helical axis. Once the helical parameters are determined, imposing helical symmetry during SPA enables 3D helical reconstruction.

However, helical reconstruction is still not a trivial task, as it requires accurate determination of the helical symmetry parameters, which are often unknown or variable. The process of determining the helical symmetry parameters called “indexing”, which is an essential and difficult step for helical reconstruction, as the helical parameters play a very important role in the alignment and averaging of particles during reconstruction. Blindly applying incorrect helical symmetry will lead to misalignment of the particles, thereby obtaining an incorrect structure. The helical parameters can be composed of two parameters: the shift distance between two neighbor assembling units named rise; and the rotation angles between two neighbor assembling units defined as twist. The traditional method for indexing is based on Fourier-Bessel analysis, which involves identifying the layer lines in the 2D power spectra of the helical projections and assigning them with integer indices that correspond to their Bessel orders. And the key step of it is to find a pair of basic vectors in the diffraction space, taking the diffraction center as the origin, so that all the diffraction points in the diffraction space could be represented by a linear combination of these vectors (Hawkes and Valdré, 1991; Toyoshima, 2000; Ward and Moody et al., 2003; Zhang and Sun, 2013; Egelman, 2015). The helical parameters can then be calculated from the spacing of the layer lines and the shift distance of the center of diffraction by using the coordinate of basic vectors (Hawkes and Valdré, 1991; Diaz and Rice et al., 2010; Egelman, 2010). Based on this principle, some programs have been developed to assist users in the indexing of diffraction pattern, such as the GUI program by YoneKura (Yonekura and Toyoshima et al., 2003), Windex (Ward and Moody et al., 2003), an online tools called Helixexplorer (https://rico.ibs.fr/helixplorer/helixplorer) and PyHI (Zhang, 2022). But these methods need to try the basic vectors manually. For users who do not have enough indexing experience, there may be omissions in this process, resulting in the failure to get the correct solution. Sun and Gonzalez et al. developed a program named HI3D (Sun and Gonzalez et al., 2022), which is based on least squares fitting between images and projections in real space. But HI3D needs high-quality asymmetry 3D map serving as prior which is usually not available. Recently, Mingtao Huang et al. combined cryo-electron tomography (cryo-ET) with a cylindrical unwrapping approach to directly calculate helical parameters(Huang, 2025). This method facilitates user understanding of the parameter derivation, but the accuracy of the calculations is compromised by cryo-ET’s extremely low signal-to-noise ratio and missing wedge artifacts. Additionally, for dynamically assembling samples, reliable parameter-based classification remains unachievable. Daoyi Li et al, based on the idea of linear regression, presented Helicon, a tool for helical parameter indexing directly by using Cryo-EM 2D images in real space, and the robustness of the method is improved through imaging stitching and L1 regularization(Li and Zhang et al., 2025). This method performs well in indexing the helical parameters of amyloid protein fibers. However, due to the dense search strategy adopted by this method, it will face the problem of large computational load. And there are still difficulties in the process of handling dynamic helical assemblies with variable diameters.

Helical reconstruction module also included in current popular software RELION (He and Scheres, 2017; Cook and Manka et al., 2020; Scheres, 2020; Thurber and Yin et al., 2021), CryoSPARC (Punjani and Rubinstein et al., 2017) and byopt-helix(Ohashi and Maeda et al., 2023), but there is no module for helical parameter calculation available. Due to the lack of a theoretical basis for parameter calculation, these methods often can only perform exhaustive searches within an extremely broad guessing range. This leads to aimlessness and uncertainty in structural computations, as well as a waste of computational time and resources. In more severe cases, it may even result in getting trapped in locally optimal but incorrect structures.

In view of the limitations of existing methods for 3D helical reconstruction, we have developed a full-process automated software, AI-HEAD, which includes automatic filament or tube diameter classification,automated diffraction indexing,automated and rational calculation of helical parameters, semi-automatic parameter sorting,the initial establishment of accurate 3D helical density maps and high resolution structure refinement (Figure 1). AI-HEAD offers the following advantages: Firstly,it can provide reliable calculations of candidate parameters based on information in Fourier space, avoiding unfounded attempts and the computational time consumption associated with large-range searches. Additionally, it eliminates the risk of exhaustive searches falling into local optima or incorrect solutions. Secondly, it can perform helical assembly simulations for candidate parameters and compare the simulated diffraction patterns with real diffraction images, thereby screening out patterns where the simulated and real diffraction match, achieving the goal of parameter sorting. Another exciting feature is that AI-HEAD can perform diameter pre-classification, which is crucial for the structural analysis of dynamically assembled samples. Only with accurate diameter-based classification can the precise helical parameters and correct structure of each class be determined.

**Figure. 1.**
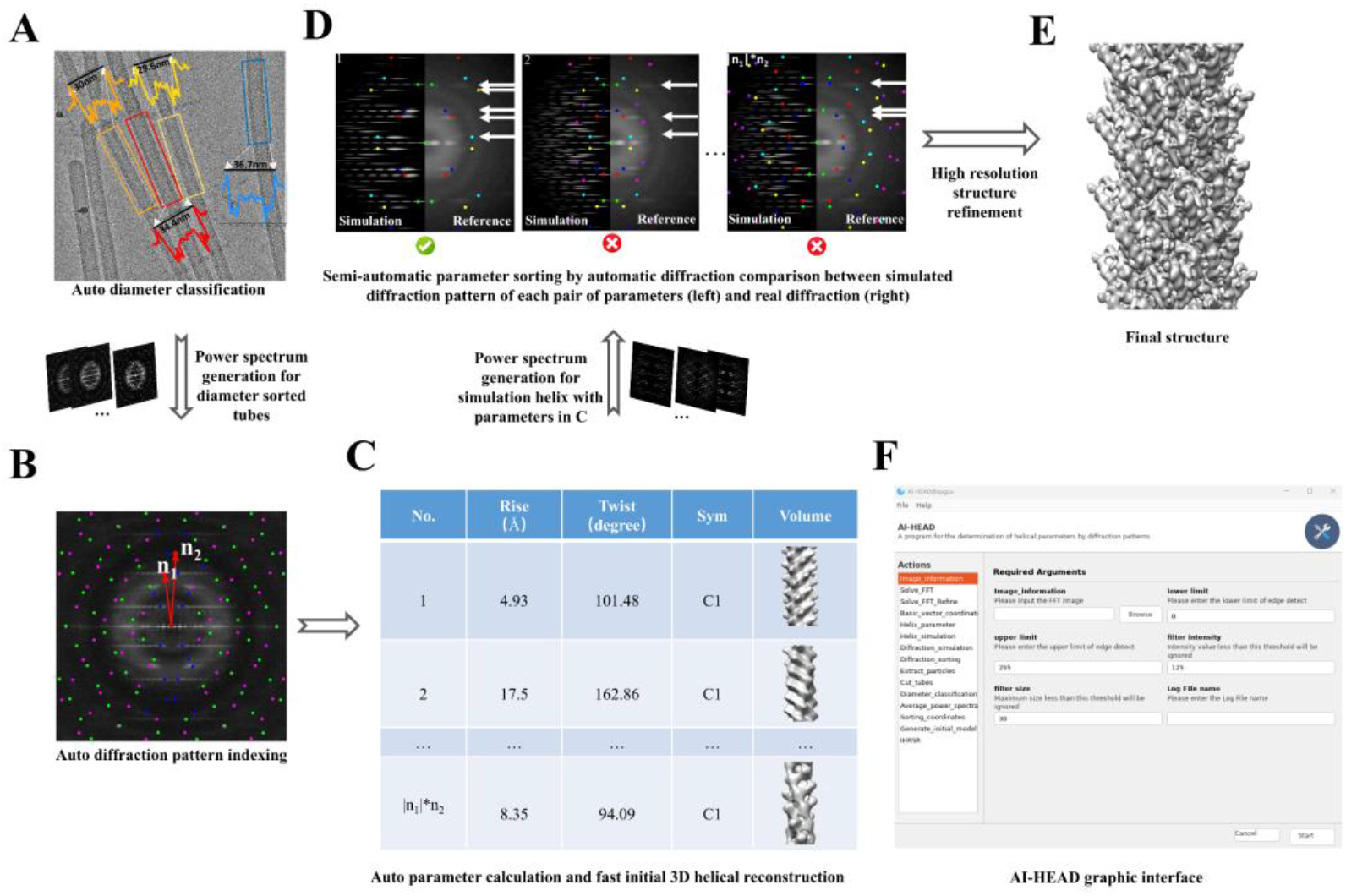
The main workflow of AI-HEAD. **A**, Diameter classification and average power spectrum generation of helical tubes. **B**, Auto-indexing of power spectrum. **C**, Calculation of helical parameters and ab-initial model. **D**, Sorting parameters by diffraction simulation. **E**, High-resolution helical refinement in CryoSPARC by using correct parameter from AI-HEAD. F, The Graphical interface of AI-HEAD.

We first validated AI-HEAD using simulated helical assemblies with ground-truth parameters as reference. The simulation experiments demonstrate that AI-HEAD can rapidly and accurately accomplish diffraction indexing and parameter computation, and efficiently identifying the correct solution by comparing with the ground truth (Extended Figure 1,Table S1 and S2).

Building upon validation with simulated ideal data, we further verified the effectiveness of AI-HEAD on real cryo-EM data. We systematically tested helical specimens with varying assembly types and data quality, including: SIRV2 (EMPIAR-10060, EMD-6310) which achieved 3.8 Å high-resolution structure and highly dynamic SNX1 complex (EMD-30592) that only reached medium resolution 9.0 Å;the thin MAVS CARD filament (EMPIAR-10031, EMD-6428) with extremely low signal-to-noise ratio and the thick TMV tube (EMPIAR-10020, EMD-2833) with well contrast.

For these four representative datasets, AI-HEAD consistently demonstrated the capability to rapidly and accurately determine the correct parameters, achieving results fully consistent with those in the EMDB database. (Extended Figure.3, Table S3-6)

Typically, the structural determination of helical assemblies faces exacerbated challenges when dealing with dynamic samples exhibiting diverse diameters - sometimes even continuous variations - which further complicates parameter acquisition. Here, we successfully determined three dynamically assembled protein-liposome helical tube complexes but with high resolution by using AI-HEAD, which are formed by respiratory syncytial virus (RSV) matrix protein (M) and large unilamellar vesicles. Through diameter classification and auto-indexing (Figure2A and 2B), helical parameters were rapidly determined from three different classes of tubes, and further refinement successfully resolved the maps to 4.2 Å, 3.7 Å and 4.6 Å resolution respectively (Figure2C and 2D). This resolution can support atomic model building. The tubes have different diameters but show similar assemble pattern as RSV M dimers stay as building blocks (Figure2C). There is no significant difference in inner-rung interactions between adjacent building blocks. Nevertheless, interactions between helical strands are completely different among three lattices, indicating inter-rung slide happened during tube formation, which results in varied tube diameters. This is consistent with the diameter variability of the RSV virions observed by using cryo-EM (Liljeroos and Krzyzaniak et al., 2013; Ke and Dillard et al., 2018). The application shows that AI-HEAD is able to classify tubes and screen for precise helical parameters accurately and rapidly, even for dynamic samples. This can promote the structure determination of membrane containing flexible helical samples, and facilitate the interpretation of membrane deformation mechanisms.

**Figure. 2.**
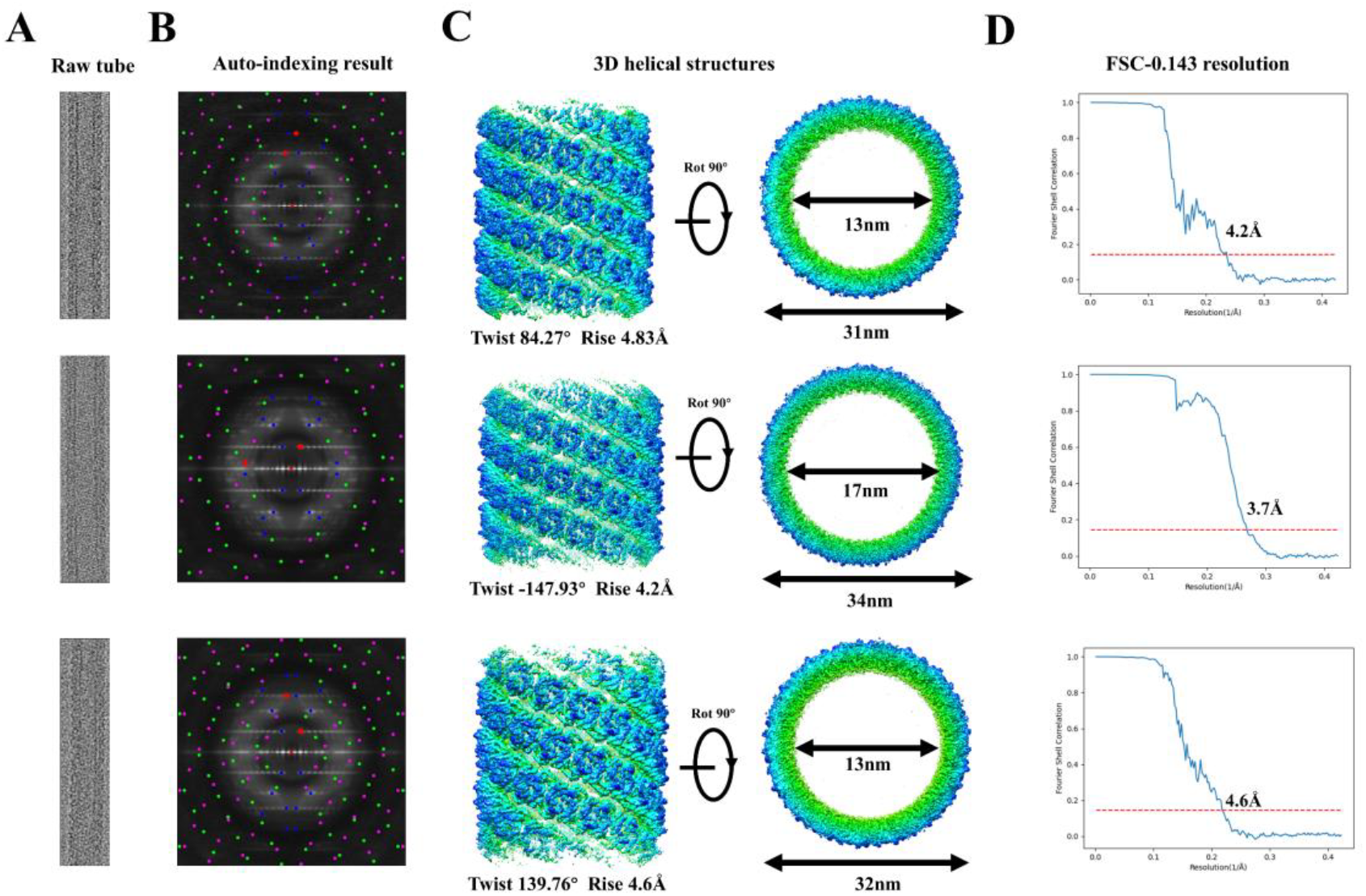
Structure determination of different RSV M helical assemblies on liposomes. **A**, Raw images of RSV M-liposome helical tube complex in different diameters. **B**, The results of the average power spectrum and Auto-indexing. **C**, High-resolution maps of different RSV M helical assemblies on liposomes. **D**, The FSC curves of the structures, with 0.143 as the cut-off standard.

## Discussion

For any helical structure, Fourier-Bessel analysis is always the most versatile and efficient way to determine the helical parameters. For researchers with no indexing experience, manual indexing is often time-consuming and laborious, and often introduces errors and leads to wrong results. AI-HEAD takes the 2D average power spectra from diffraction patterns as input to achieve automatic indexing and outputs the corresponding parameters, significantly reducing the time and complexity of the indexing process.

However, the program still has some limitations that affect its performance and accuracy. One major challenge is that, although the program can perform a simple sorting of results by simulation, the only correct helical parameters cannot be determined as there are still many candidate parameters that cannot be sorted. This means that the correct parameters must be obtained by conducting helical reconstruction based on each possible set of helical parameters, which increases the overall time required for data processing. To solve this problem, AI-HEAD provides the interface of SPIDER/IHRSR, which could automatically perform helical reconstruction in batch based on the remaining helical parameters. And to increase the speed of processing, particles could be down sampled before reconstruction (e.g., bin4, bin2). The helical parameters could be firstly tested using these down sampled particles, and once the correct parameters are identified, the particles can be gradually restored to their original size for high resolution refinement.

Another challenge is the sensitivity of the algorithm to noise and outliers, which can cause failures in properly identifying and segmenting diffraction patches. This may lead to increased manual corrections and introduce errors into the indexing results. In fact, the quality of the power spectra is strongly related to the success of the indexing process. A power spectrum which contain less high resolution information may have no unique solution at all due to factors such as out-of-plane tilt, sample heterogeneity, and noise in cryo-EM data. To overcome these issues, the homogeneity of the sample should be ensured as much as possible during sample preparation. If the helical assembly of the sample is highly dynamic, the data could be pre-classified appropriately before processing. Indexing can then be performed on one selected class, and the calculated parameters from this class can be used to carry out the subsequent helical reconstruction.

In the future, we’ll propose some more improvements and enhancements to the existing program. First, we plan to introduce neural networks to train with 2D power spectra of real Cryo-EM data, so as to identify diffraction regions more accurately instead of edge detection, thus reducing manual intervention. Additionally, we plan to develop a faster alignment algorithm based on the IHRSR algorithm, enabling the program to achieve rapid parameter screening after indexing. Thus, it is possible to input particles and their power spectra into our programs to obtain the corresponding structure in a short time. With that, the helical reconstruction problem can be automated to a greater extent, and the helical assembly structures of various paradigms can be analyzed efficiently and in batches, which will provide strong technical support for exploring biological issues such as physiology, pathology and membrane dynamics.

## Methods

### Protein preparation and data collection

RSV M protein was expressed as His6 fusion protein in a pET28a vector in Escherichia coli BL21(DE3). Cells were grown from fresh starter cultures in LB broth for 3.5 h at 37°C, followed by induction with 0.5 mM IPTG at 16°C overnight. Cells were lysed by sonication in lysis buffer (50 mM NaH2PO4-Na2HPO4, 300 mM NaCl, pH 7.4) plus lysozyme (1mg/ml, BBI), protease inhibitors (Beyotime), RNase (12 μg/ml, Sigma), and 0.25% CHAPS {3-[(3-cholamidopropyl)-dimethylammonio]-1-propanesulfonate} (Beyotime). Lysate were clarified by centrifugation and the soluble His6-M protein was purified by using Nickel Sepharose 6 Fast Flow resin (Cytiva). The bound protein was washed with lysis buffer plus 50 mM imidazole and incubated with 3C protease overnight at 4°C to remove the His6 tag. The processed protein was concentrated and further purified on a Superdex 200 Increase 10/300 GL column (Cytiva) in lysis buffer.

Large unilamellar vesicles were prepared by combining POPC (1-palmitoyl-2-oleoyl-glycero-3-phosphocholine), POPE (1-palmitoyl-2-oleoyl-snglycero-3-phosphoethanolamine) and POPS (1-palmitoyl-2-oleoyl-sn-glycero-3-phospho-L-serine) in a ratio of 2:5:3, and dried under a steady stream of N2 (all lipids were purchased from Avanti Polar Lipids, Inc.). lipid films were hydrated in binding buffer (25 mM Tris-HCl, 150 mM NaCl, pH 7.4) using freeze-thaw cycles. RSV M protein was then incubated with liposomes for 4 h at room temperature to form M-liposome helical tube complex.

3 μl of M-liposome complex was transferred onto 300 mesh quantifoil holy carbon R1.2/1.3 gold grids which were glow-discharged before use. Excess solution was blotted for 3.5 s in a Vitrobot Mark IV (Thermo Fisher Scientific) at 4 °C and 100 % humidity before grids were rapidly plunged into liquid ethane. Frozen grids were loaded into a Titan Krios G4 (Thermo Fisher Scientific) operated at 300 kV, equipped with a Selectris-X energy filter (10 eV slit width) and Falcon 4i direct electron detector (Thermo Fisher Scientific). Images were taken with a pixel size of 1.18 Å/pixel at 1 to 2 μm underfocus in counting mode by using EPU2, and a total dose of 40 e/Å^2^ was applied.

### Diameter classification of helical assemblies

AI-HEAD could extract all the tubes by “Extract_particles” module based on coordinates in EMAN2 format. All the tubes are projected along the helical axis to get the 1-dimension projection curve. Subsequently, the diameters of all the tubes could be obtained by measuring the peak distance of the projection curve and classified into multiple classes according to the specified number of classes. In addition, the tube coordinates of a specific diameter could be obtained by sorting the original tube coordinates through the “Sorting_coordinates” module of AI-HEAD.

### The attributes of basic vectors

The ideal continuous helix is the result of periodic extension of helical density. At this point, the only parameter of the helix is the period of the helix (pitch). According to Fourier-Bessel analysis, a continuous helix in Fourier space is equivalent to sampling in the reciprocal of the pitch. In this way, a “diffraction layer line” is formed in Fourier space, and the intensity of each diffraction layer line is proportional to the square of the Bessel function corresponding to the Bessel order (Hawkes and Valdré, 1991; Ward and Moody et al., 2003; Diaz and Rice et al., 2010; Egelman, 2010; Zhang, 2013). The Bessel order of each diffraction layer line could be estimated by the location of the maximum point on the layer line (See “Basic_vector_cooordinate” in supplementary data in detail). Therefore, in the case of a continuous helix, we only need to find the basic vector 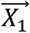 to obtain the pitch of helix at this time (Extended Figure.1A).

Helical assemblies are formed by assembling biological macromolecules in a helical arrangement, which could be determined by the pitch of the helix and the rise distance between two neighbor assembling units which corresponding to one single rotation of the asymmetric unit. Therefore, a helical assembly could be seen as the result of sampling the ideal continuous helix in real space with equal distance, and this distance is helical rise. Equivalently, the diffractions pattern of continuous helix in the Fourier space was extended with the reciprocal of helical rise. To get the rise of helical assembly, we need to find the first point on the meridian above the equator. To get the pitch, one has to find the equal distance between each layer line of the elementary or first diffraction of the ideal helix. (See “Helix_parameter” part in supplementary data in detail).

The mathematical relationship between the diffraction vector and the pitch and rise ensures the calculation of our parameters (see Helical_parameter session in the Supplementary Data). Therefore, the identification and indexing of the basic vectors of diffraction pattern is a prerequisite for the calculation of helical parameters. To better understand the selection principles of the basic vectors in diffraction space, we first verify the equivalence between vectors in diffraction space. Here, we introduce a pair of vectors 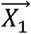 and 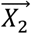 as basic vectors, so that all the diffraction points on the diffraction pattern could be represented by a linear combination of them with integer coefficients. (Extended Figure.1B). In this way, helical parameters could be obtained through the linear combination of the basic vectors and corresponding reciprocal space transformation calculation.

Each pair of basic vectors can generate the whole diffraction lattice. We define the basic vectors that produce the same lattice “equivalent basic vectors”. In the following, we will introduce the conditions for two pairs of basic vectors to be equivalent to each other. Let’s assume that one lattice is generated from 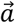 and 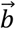 as the basic vectors, and can also be generated by 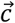 and 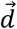. As lattice points are generated by a linear combination of the basic vectors with integer coefficients, 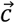 and 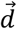 must be represented by linear combination of 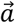 and 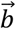 with integer coefficients.

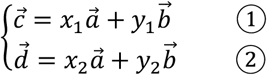

In the same way, 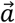 and 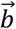 must also be represented by a linear combination of 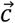 and 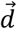 with integer coefficients.

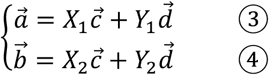

By combining the equations ➀, ➁, and ➂, the following equations are obtained:

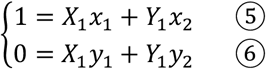

Then *X*_1_ and *Y*_1_ can be solved by combining equations ➄ and ➅:

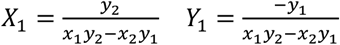

By combining the equations ➀, ➁, and ➃, the following equations are obtained:

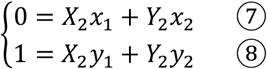

And *X*_2_ and *Y*_2_ can be solved by combining equations ➆ and ➇:

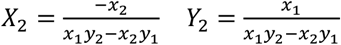

Let M represents *x*_1_*y*_2_ − *x*_2_*y*_1_. As *x*_1_, *x*_2_ *y*_1_, *y*_2_, *X*_1_, *X*_2_, *Y*_1_, *Y*_2_ are all integers, thus M is their greatest common divisor. In particularly, this condition is always true when M is equal to ±1. From this we could infer that the lattice generated by 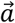 and 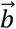 could also be generated by 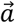 and 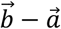 (In this case *x*_1_ = 1, *y*_1_ = 0, *x*_2_ = −1, *y*_2_ = 1).

**Extended Figure. 1.**
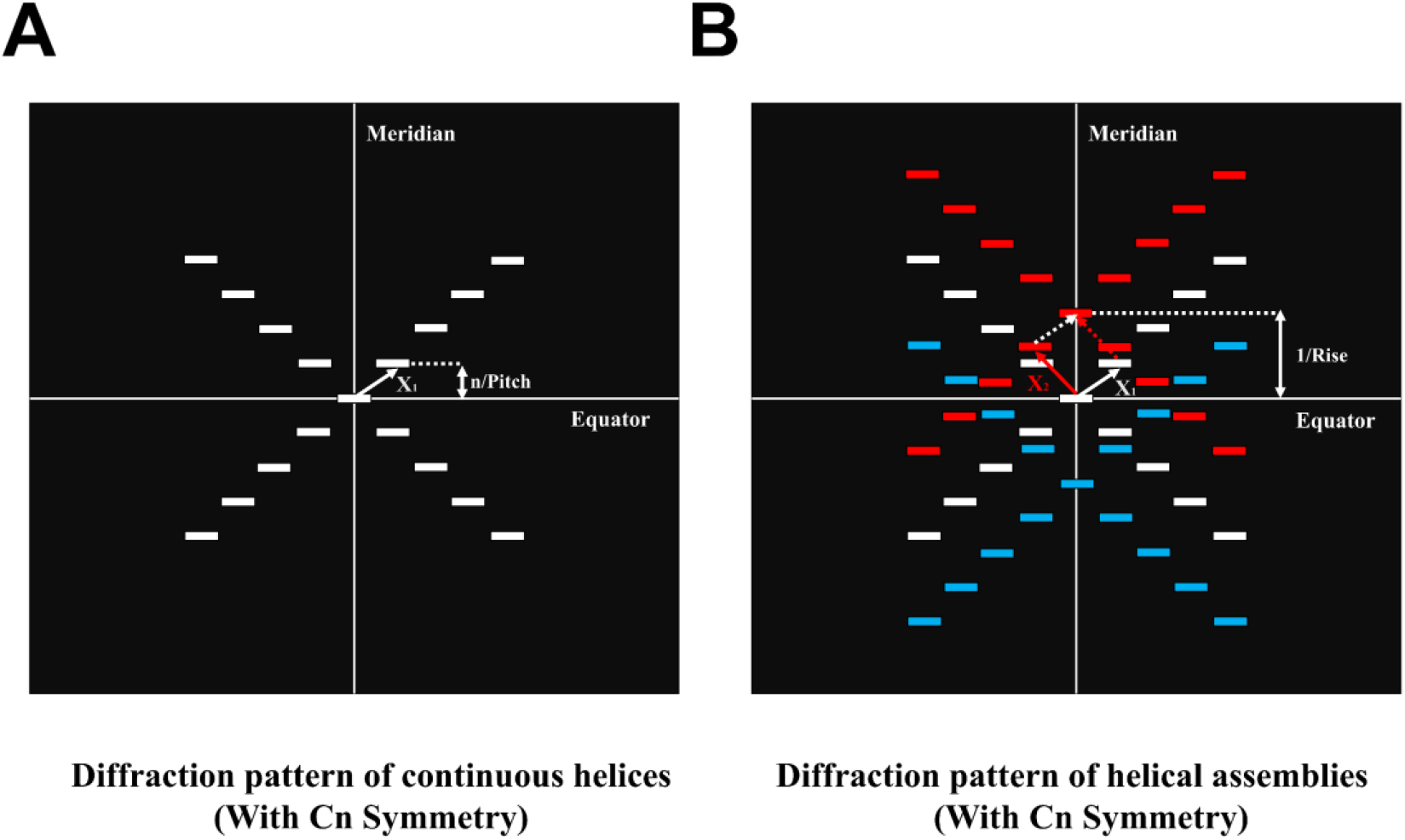
Diffraction patterns of continuous helices and helical assemblies. **(A)** Diffraction pattern of continuous helices. The diffraction points are arranged on equally spaced diffraction layers, and the distance between the layers is the reciprocal of the period of the helices (Pitch). **(B)** Diffraction pattern of helical assemblies. The helical assemblies are equivalent to an equidistant sampling of the continuous helices in the unit of rise, which in Fourier space is equivalent to an equidistant convolution continuation in the unit of the reciprocal of rise. All other diffraction points can then be represented by linear combinations of 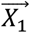 and 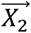 with integer coefficients.

The process of AI-HEAD program to obtain helical parameters could be divided into 4 parts: Auto-indexing, Calculation, Auto-simulation and Diffraction-sorting. In the “Auto-indexing” part, we determined the position of the basic vectors; And then we use the information of basic vectors in Fourier space to calculate a series of possible parameters in “Calculation” part; Next, in order to perform a simple filtering of the calculated parameters, we simulated the projection and the diffraction process of the helices in “Auto-simulation” part which is based on the helical parameters calculated before; With that, we could sort the results by comparing the diffraction pattern of simulation with the diffraction pattern of the original data in “Diffraction-sorting” part. Each part could be achieved by several different modules. In the following sections, we will sequentially introduce these four interlocking steps for parameters determination.

### Auto-indexing of helical diffraction pattern

First of all, before indexing, what we need first is the diffraction pattern, that is, a 2D power spectrum which only contains information of the main layer lines in Fourier space. The diffraction pattern could be obtained from a single tube by adjusting the power spectrum to preserve the main diffraction signal and remove the excess noise. However, during the process of indexing, signals in low-frequency are often more reliable. Generally, the average of 2D power spectra will contain more information in low frequency, and it may be efficient to get the desired result by using it as the beginning of indexing. The diffraction pattern could be obtained by the “Basic_vector_coordinate” module of AI-HEAD, which could not only obtain the power spectrum from raw data, but also conduct adjustment of power spectrum and save it. Once the diffraction pattern is obtained, indexing can be carried out next.

As mentioned earlier, the key step of indexing is to find a pair of basic vectors in Fourier space (Hawkes and Valdré, 1991; Toyoshima, 2000; Ward and Moody et al., 2003). Due to the influence of helical curvature, diffraction points in Fourier space are modulated by Bessel function into patches, and these patches are arranged on a set of lines, which is called “layer lines”. From the above properties, it can be deduced that there exists a pair of basics vectors starting from the diffraction center, and the lattice generated by them can cover most of the diffraction patches in Fourier space. According to this property, we used “Global searching” strategy in AI-HEAD. We sort the basic vectors based on the matching results between the lattice generated by them and diffraction patterns by traversing all the candidate basic vectors, thereby automatically selecting the more ideal basic vectors. First, all the positions of the diffraction patches in the diffraction pattern could be marked by the “Image_information” module of AI-HEAD. This module will use Canny algorithm (Canny, 1986) to automatically detect the edge of the diffraction pattern and perform the threshold segmentation (N., 1979; A. and S. et al., 2018; D. and H. et al., 2018; Latychevskaia and Fink, 2018; N. and C., 2018; P. and K., 2018) to find out the coordinate information of all the diffraction patches in the diffraction pattern. Due to the complexity of diffraction pattern, the Canny algorithm may not be able to completely find all the diffraction patches or segment some connected diffraction patches in Fourier space. So, we could manually supplement the missing patches or manually segment the incomplete fully automatic diffraction patches to mark all the regions for indexing. In addition, AI-HEAD also use FAST algorithm (E. and R. et al., 2010) to automatically detect all the feature points of the diffraction pattern, which could be used later to find the basic vectors.

Following the annotation of the diffraction pattern, the central coordinates of the pattern and the positions of all diffraction patches were determined. Subsequently, within the “Solve_FFT” module, each diffraction patch is condensed into a discrete diffraction point in Fourier space. Two such points are then selected sequentially to form a pair of basis vectors with the diffraction center. Using these basis vectors, a series of lattices and their symmetric counterparts can be generated, collectively forming an extensive solution set. The Fourier space of the diffraction pattern is divided into four quadrants using the “equator” and “meridian” as boundary lines. Ideally, helical diffraction exhibits high symmetry with respect to both the equator and meridian (Toyoshima, 2000). Leveraging this symmetry, the search space for the AI-HEAD algorithm is significantly reduced, requiring only one representative point from the first and second quadrants, respectively, to form the necessary vector pairs with the diffraction center. Even so, the “solution set” of basic vectors is very large because it records all possible combinations of basic vectors.

Once the “solution set” is determined, AI-HEAD will evaluate each of the solutions in order and then rank them. As mentioned above, under the ideal indexing, the lattice generated by the basic vectors should have a good match with the original diffraction pattern. When the diffraction pattern contains more information in high resolution, it can be observed that some of the diffraction patches are located on the meridian. According to the principle of the Fourier-Bessel transformation, the vertical distance between the diffraction patch located on the meridian and the diffraction center is proportional to the reciprocal of the rise of the helical structure (Hawkes and Valdré, 1991). Therefore, when there are diffraction patches on the meridian, the lattice generated by the basic vectors must be exactly all on the diffraction patch near the position of the meridian. If the lattice does not completely cover the diffraction patch on the meridian, or covers the area without diffraction patch on the meridian, then this set of basic vectors is not ideal. In addition, the lattice generated by the ideal basic vectors should be distributed on more diffraction layer lines, and on this basis cover more diffraction points. AI-HEAD will automatically sort the solution set according to several conditions of the ideal indexing, and we could specify the top n groups of indexing solutions to output. In general, the top 30-50 solutions should contain the ideal indexing results, and in the case of fewer diffraction patches, a smaller number can be taken for output. Each output diffraction pattern is marked with points of different colors and size: the diffraction center and the diffraction points which formed the basic vectors with diffraction center are shown in red (For the purpose of differentiation, the size of the points of the two basic vectors is larger than the diffraction center); The lattice points generated by the basic vectors are shown in pink; Symmetrical lattice points which is symmetric with the lattice about the meridian are shown in light green; Lattice points that fall exactly on the diffraction point are shown in blue. Then the most ideal solution could be chosen to get the relative position of basic vectors and then to conduct the calculation of helical parameters.

As the “Global searching” strategy abstracts all the diffraction patches into diffraction points by using the center of diffraction patches, sometimes the center of patches not the best position to form a basic vector. As mentioned earlier, for one diffraction patch, FAST algorithm could find plenty of feature points. “Solve_FFT” module provides the user with the selection of basic vectors from a global perspective, and tell the user the optimal diffraction patches. On the basis of that, we could select the diffraction patches region by the “Solve_FFT_Refine” module, and then find the precise position of basic vectors in the diffraction patch. Sometime this step is not necessary, but it may work well and provide more solutions when the diffraction pattern is noisy, or the diffraction patches are very long. At this point, the “Auto-indexing” part has been finished.

### Calculation of helical parameters

Once the ideal basic vectors are determined, the helical parameters could be calculated based on the coordinates of these basic vectors in Fourier space (see “Helical_parameter” in supplementary data in detail). AI-HEAD could automatically transform the basic vectors coordinates in Fourier space to the n-Z coordinates based on the radius of helical assembly according to Fourier-Bessel transformation (Hawkes and Valdré, 1991; Diaz and Rice et al., 2010) for subsequent calculation. Sometimes, one of the vectors chosen may point at diffraction patches which lie on the “meridian” whose Bessel order is zero, which could not be used to calculated the helical parameters directly (Extended Figure.2D(a)). Since the same lattice could be generated by various vectors which satisfy certain conditions (see “The attributes of basic vectors” session in method), the zero Bessel order vector pair could be converted into another equivalent vector pair by linear combination (Extended Figure.2D) for parameter calculation. As the radius of the helical assemblies may not be very precise, and the diffraction points is modulated by the Bessel function, the Bessel order of the vectors would become a range (Hawkes and Valdré, 1991). With that, we could easily obtain a set of candidate helical parameters by “Helix_parameter” module of AI-HEAD. And now, the “Calculation” part has been finished.

### Auto-simulation of projections and diffraction based on different parameters

Although there are many groups of possible helical parameters calculated in the previous part, the corresponding parameters of each helical assembly should be unique, so the subsequent parameter screening needs to be carried out by helical reconstruction based on these parameters in turn using IHRSR. However, when the radius of the helical assembly is relatively large (for example, 200Å), the number of possible results is also very large, which will undoubtedly increase the time cost of helical reconstruction. Some of the parameters calculated in previous step are very close to each other, and some are very different, which means there will be diffraction patterns of helical projections based on some parameters, which are very different from the diffraction pattern derived from raw data. In the “Auto-indexing” part, we find the approximate location of the basic vectors and the lattice generated by them based on the diffraction pattern of raw data. So, we can use it as a reference to filter out some unreasonable parameters. This process could be divided into three parts. Firstly, a set of helices projections based on the radius and the candidate helical parameters are simulated with “Helix_simulation” module of AI-HEAD. At the same time, AI-HEAD will also calculate the distance between the corresponding diffraction layer lines and the distance of the diffraction center along the meridian according to each set of helical parameters, and this information will be used to generate diffraction lattice later. Next, all of the diffraction patterns of simulated helical projections could be got by using “Diffraction_simulation” module in AI-HEAD. Once the simulated diffraction pattern and the original diffraction pattern at the same scale was obtained, the “Auto-simulation” part is complete.

### Sorting the results based on the simulated lattice and diffraction pattern

Next, we will compare the simulated data with the original data according to the diffraction pattern and the simulated lattice, which could be achieved by using “Diffraction_sorting” module. AI-HEAD will automatically match half of the simulated diffraction pattern with half of the original data diffraction pattern to form a “converge image”. In this way, we can align the layer lines of the simulated diffraction pattern with those of the raw data. Then, since the “Auto-simulation” part also generates on layer line information for each set of parameters, with that, AI-HEAD can also combine the information of layer lines to generate a simulation lattice for the corresponding parameter on each “converge image”. Based on all of the information above, we could use the lattice generated in “Auto-indexing” as a reference; to filter out the parameters whose lattice arrangement is completely different from reference. Finally, the remaining parameters can be used for helical reconstruction using IHRSR one by one to converge to get the only correct helical parameters (Video S1-3).

### Get the parameter of Simulation Helix by using AI-HEAD

In order to test the function of the AI-HEAD, we first created a simulation helix based on the given helical parameters (rise=25.6Å, twist=45°, radius=100 Å, C1) by SPIDER, and use AI-HEAD to conduct the indexing and the calculation of helical parameters (Extended Figure.2). We projected the simulation helix to obtain its projection. Then, we conduct the 2D Fourier transformation and get the power spectrum of the projection. Next, we adjust the contrast and brightness of the power spectrum to keep the main diffraction patches and thus get the diffraction pattern. These steps could be finished by using Basic_vector_coodinate module to display and adjust the power spectrum in AI-HEAD. In this case, we choose top3 results which have the same lattice but generated by different basic vector pairs. Then we get the coordinate of the basic vectors in turn by using Basic_vector_coodinate module, and input them into the “Helix_parameter” module to calculate the helical parameters. After calculation, all of the solution set contained the correct parameters (See Table.S1-2 in detail). It proves that the AI-HEAD could accurately obtain the helical parameters of the helical structures under the ideal condition of complete noise-free interference.

**Extended Figure. 2.**
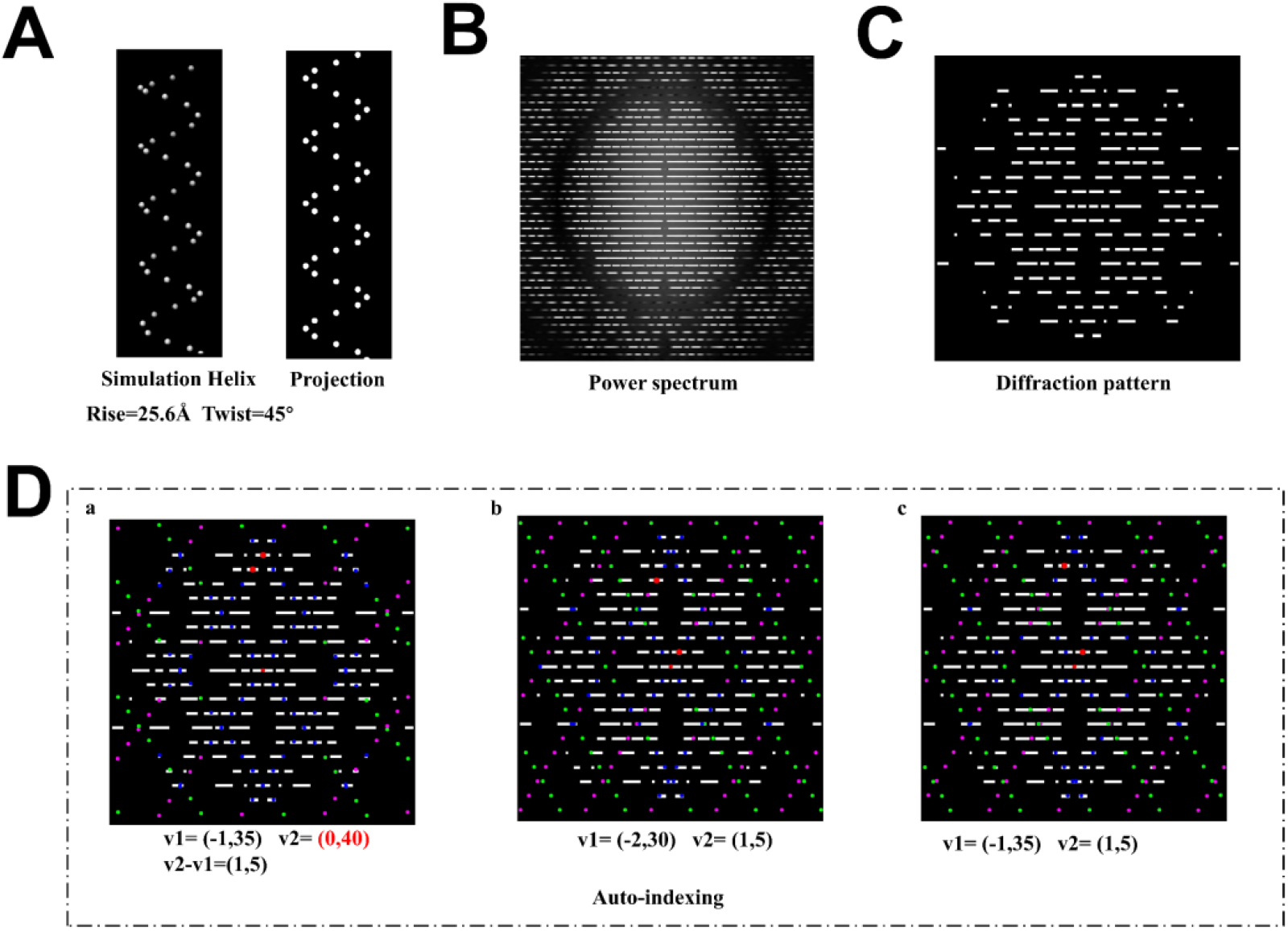
The Auto-indexing results of simulation data. **(A)** The simulated 3D helix was generated by SPIDER with definite helical parameters, and its projection could be conducted by using EMAN2. **(B)** The power spectrum of the projection. **(C)** The diffraction pattern obtained by power spectrum after adjusting. **(D)** Several ideal results of auto-indexing: Red points(small): the center of diffraction pattern; Red points(large): the basic vector pair; Pink points: the lattice generated by the basic vector pair; Green points: the symmetric lattice generated by the basic vector pair; Blue points: the points of lattices which covered the diffraction patches.

### Get the parameters of helical assemblies from EMDB by using AI-HEAD

Then, we further tested the performance of AI-HEAD on real cryo-EM data. We chose several helical structures which have different resolutions (MAVS CARD (EMD-6428, 4.2 Å, C1), TMV (EMD-2833, 4.0 Å, C1), SIRV2 (EMD-6310, 3.8 Å, C1), SNX1 complex (EMD-30592, 9.0 Å, C1)) from EMDB. The density maps in relatively high resolution (MAVS CARD, TMV and SIRV2) corresponds to helical data which is in high quality while the map in lower resolution (SNX1 complex) corresponds to the relatively poor-quality helical data. To test the performance of AI-HEAD to calculate helical parameters, we started the indexing process with each set of raw data. For MAVS CARD (EMPIAR-10031) and TMV (EMPIAR-10020), there are complete micrographs and particle coordinates data in EMPIAR; For SNX1, We used the micrographs and particle coordinates data from our previous work (Zhang and Pang et al., 2021). So, for these three data sets, we first conduct preprocessing in Relion and extract the particles, then import them into CryoSPARC. After 2D classification in CryoSPARC, we select the best classes to obtain the average power spectra by using “Average power spectra” module in CryoSPARC. For SIRV2(EMPIAR-10060), there is only particle set data with SPIDER format in EMPIAR. Therefore, we directly used the particle set to generate the average power spectrum by SPIDER. Since longer helices tend to produce more sharp diffraction layer lines, before obtaining the average power spectrum, we padded the particles in real space until the box size reached approximately 2000. We then conduct auto-indexing and calculation of helical parameters by using AI-HEAD. (See Extended Figure.3, Table.S3-S6). This proves that AI-HEAD can not only solve the helical parameters and perform high resolution reconstruction for good quantitative data, but also obtain the correct solution for poor quantitative or dynamics data.

**Extended Figure. 3.**
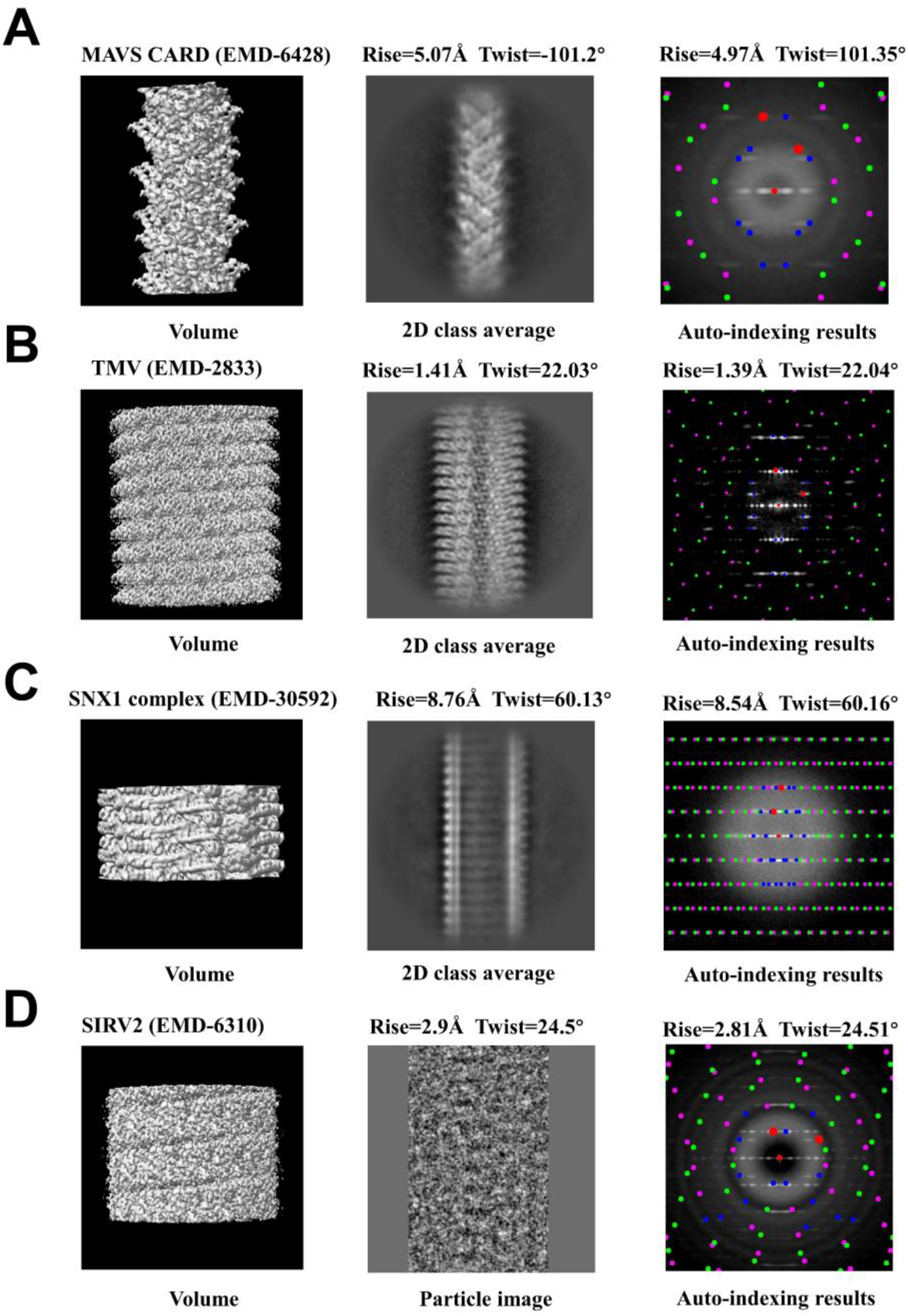
The test results of helical data from EMDB. **(A)** MAVS CRAD (EMD-6428) (parameter in EMDB: Rise=5.07Å Twist=-101.2°; Calculated by AI-HEAD: Rise=4.95Å Twist=101.2°). **(B)** TMV(EMD-2833) (parameter in EMDB: Rise=1.41Å Twist=22.03°; Calculated by AI-HEAD: Rise=1.38Å Twist=22.01°). **(C)** SNX1 complex (EMD-30592) (parameter in EMDB: Rise=8.76Å Twist=60.13°; Calculated by AI-HEAD: Rise=8.66Å Twist=60°). **(D)** SIRV2 (EMD-6310) (parameter in EMDB: Rise=2.9Å Twist=24.5°; Calculated by AI-HEAD: Rise=2.87Å Twist=24.55°).

### Auto-simulation and Parameter sorting

As mentioned earlier, the AI-HEAD computes a range of possible solutions based on the basic vector coordinates. When the position of the selected basic vector pair is distant from the meridian or the helix has a large radius, the resulting solution set may become substantially large. Therefore, AI-HEAD provides a way to enable a simple sorting of solution set by simulation. Now, we take the SIRV2 dataset as an example to demonstrate the simulated diffraction of AI-HEAD and the function of diffraction sorting. Since we have found the basic vectors of the diffraction pattern of SIRV2 by “Auto-indexing” before, and generated the corresponding helical lattice (Extended Figure.4A). To conduct the sorting of data, we generated the diffraction pattern of all simulated data and the lattice under the corresponding parameters, and use the diffraction pattern of the original data and the helical lattice as a reference for data sorting. Since the results of all calculations are based on the basic vectors selected in “Auto-indexing”, the simulation lattice corresponding to the correct parameters should be similar to the lattice of “Auto-indexing” (Extended Figure.4B).. However, the diffraction pattern and simulation lattice of different helical parameters in the low-resolution region in Fourier space could be extremely similar. Therefore, this approach does not strictly obtain the only correct set of parameters. And when the rise or pitch of one helix is very small, the algorithm may not be able to generate a complete simulation lattice, resulting in a simulation lattice that is very similar to a part of “Auto-indexing” result. So, to be conservative, we could keep all results that are similar to “Auto-indexing” (Extended Figure.4B, Video.S1-2) and remove results that are not similar at all (Extended Figure.4(C-D), Video.S3). In this case, 18 sets of 34 solutions were removed, which reduced the difficulty of subsequent parameter screening.

**Extended Figure. 4.**
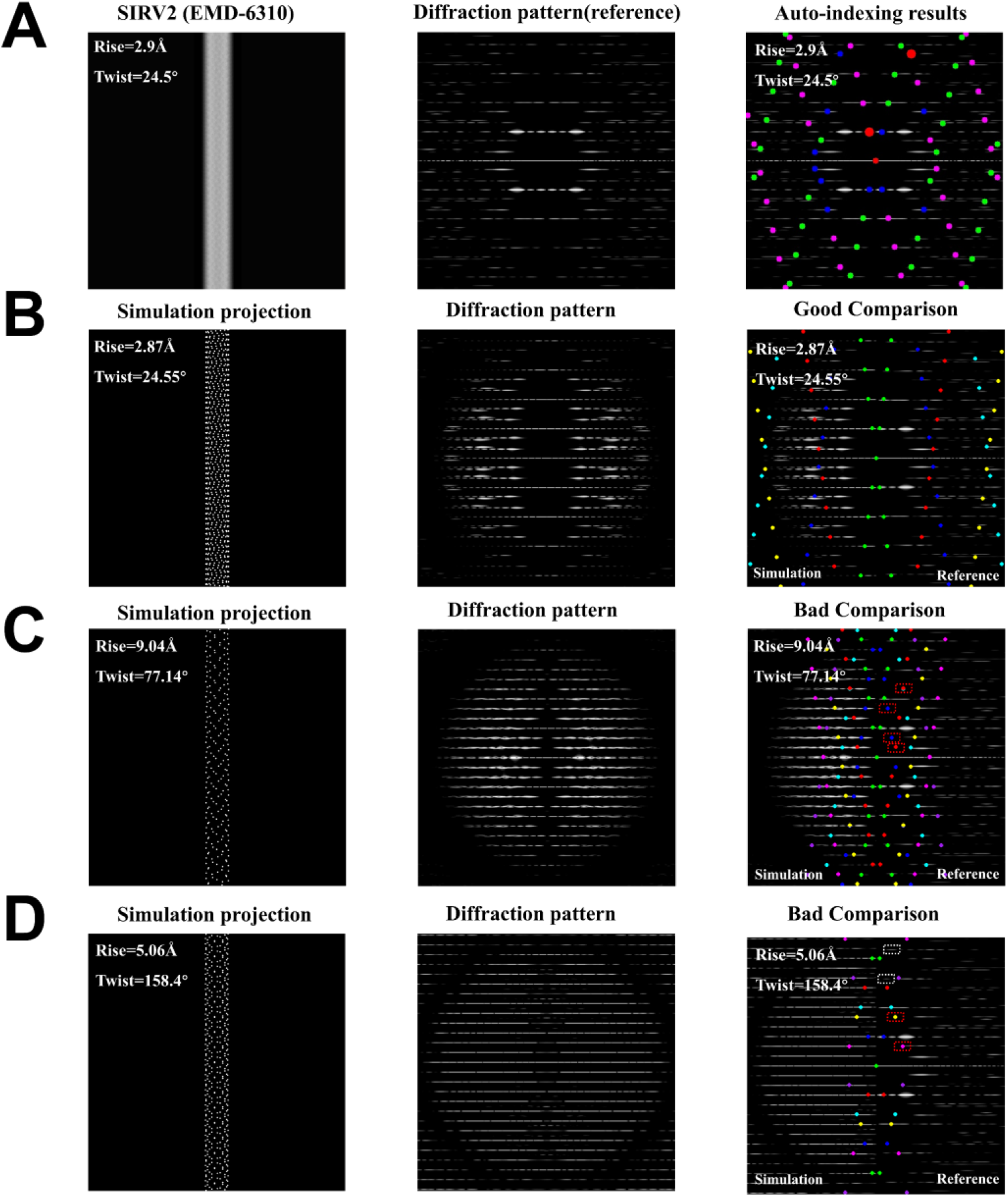
The diffraction simulation and sorting of SIRV2 from EMDB. (A) The projection, diffraction pattern and auto-indexing results of SIRV2 (EMD-6310, parameter in EMDB: Rise=2.9Å Twist=24.5°). (Points of different colors distinguish basic vectors and other points in lattice (Red points(small): the center of diffraction pattern; Red points(large): the basic vector pair; Pink points: the lattice generated by the basic vector pair; Green points: the symmetric lattice generated by the basic vector pair; Blue points: the points of lattices which covered the diffraction patches). (B-D) are the simulated helical projections generated by AI-HEAD according to the specified parameters, the simulated lattice (Points of different colors to distinguish lattice points from different diffraction centers, Purple: center located on (0, 3/rise), Cyan: center located on (0, 2/rise), Blue: center located on (0, 1/rise), Green: center located on (0,0), Red: center located on (0, −1/rise), Yellow: center located on (0, −2/rise), Pink: center located on (0, −3/rise)) generated according to the corresponding parameters, and the comparison with the real diffraction pattern(left: diffraction pattern of simulation, right: diffraction of original data): (B) The projection of simulation helix generated by AI-HEAD based parameter calculated by AI-HEAD: Rise=2.87Å Twist=24.55°. (C) The projection of simulation helix generated by AI-HEAD based parameter calculated by AI-HEAD: Rise=9.04Å Twist=77.14°. This result is significantly different from the lattice generated by Auto-indexing, where the points that do not appear in Auto-indexing are circled in red dashed boxes. (D) The projection of simulation helix generated by AI-HEAD based parameter calculated by AI-HEAD: Rise=5.06Å Twist=158.4°. This result is significantly different from the lattice generated by Auto-indexing, where the points that do not appear in Auto-indexing are circled in red dashed boxes; And the points missed compared with Auto-indexing are circled in white dashed boxes.

### Get the parameters of RSV M assemblies by using AI-HEAD

To get the structure of RSV M assemblies, we first employed the EMAN2 utility e2helixboxer.py to manually select viral tubes from micrographs. The extracted coordinates were then imported into the AI-HEAD to extract all identified virus tubes. These tubes were subsequently pre-classified within AI-HEAD based on their diameters, yielding distinct structural classes. Among these, we selected three diameter classes that contained relatively large numbers of particles: one class with a diameter of 31 nm, another with a diameter of 32 nm, and a third with a diameter of 34 nm. Using AI-HEAD, we filtered the coordinates to separate tube into 3 classes by their diameters. With these curated datasets, we computed the average power spectra for each class using AI-HEAD.

Auto-indexing of the class-specific power spectra was then performed by AI-HEAD, yielding candidate helical parameters. To refine and validate these parameters, we first applied CTF pre-multiplication to all micrographs using AI-HEAD. The pre-multiplied images were then re-extracted based on the previously defined tube coordinates and segmented into particles. These particle stacks were converted into SPIDER format (.spi) using EMAN2 and down sampled by a factor of four to facilitate efficient parameter screening.

Initial models for each diameter class were generated in SPIDER as featureless cylinders. These models, along with the candidate helical parameters, were used to batch-generate IHRSR projects within AI-HEAD. Upon executing all IHRSR reconstructions, we identified a single correct density map (Bin4) and refined helical parameters corresponding to each diameter class. The preliminary parameters identified were: for the 31 nm class, a twist of 84.24° and a rise of 4.85 Å; for the 32 nm class, a twist of 139.78° and a rise of 4.66 Å; and for the 34 nm class, a twist of –147.93° and a rise of 4.23 Å.

To improve the quality of initial reference, we utilized bin2 particles and performed further IHRSR refinement, resulting in more structurally detailed initial models and optimized helical parameters.

For high-resolution reconstruction, we re-extracted particles using the AI-HEAD-filtered coordinates and processed them in RELION for 2D classification. Well-defined classes were selected, and the refined IHRSR models served as initial references. Using the IHRSR-derived Helix_refine parameters, we obtained high-resolution reconstructions in CryoSPARC, achieving final resolutions of 4.2 Å for the 31 nm class, 4.6 Å for the 32 nm class, and 4.0 Å for the 34 nm class. Subsequent CTF refinement in RELION further improved the final resolutions to 4.2 Å, 4.6 Å, and 3.7 Å for the 31 nm, 32 nm, and 34 nm classes respectively. The final parameters identified were: for the 31 nm class, a twist of 84.27° and a rise of 4.83 Å; for the 32 nm class, a twist of 139.76° and a rise of 4.60 Å; and for the 34 nm class, a twist of –147.93° and a rise of 4.20 Å.

## Supporting information

All tables, videos and descriptions about the main

## Data availability

Simulated helix data was generated by SPIDER 23.03 with the given helical parameters. The helical density maps MAVS CARD (EMD-6428), TMV (EMD-2833), SIRV2 (EMD-6310), SNX1 complex (EMD-30592) along with their parameters could be got from EMDB. MAVS CARD (EMPIAR-10031),TMV(EMPIAR-10020) and SIRV2(EMPIAR-10060) data could be download from EMPIAR; SNX1 data are from the author’s previously published papers.

The RSV M cryo-EM sample preparation and data collection were performed at the Advanced Bio-imaging Technology Platform of Guangzhou National Laboratory.

## Code availability

The program is a Python script that runs under Python version 3.10. The script depends on the following Python packages: Pytorch 2.1.0 +cu118, mrcfile 1.1.2, numpy 1.18.3, matplotlib 3.3.1, opencv-python 4.5.5.64, pandas 1.2.4, Gooey 1.0.8.1(https://github.com/chriskiehl/Gooey). These packages can be installed on modern operating systems with standard Python library management tools such as Python Package Installer (PIP; **https://pip.pypa.io/en/stable/**) and anaconda (**https://docs.conda.io/en/latest/**). The CTF module and the particle extraction module adopt the libtilt project (https://github.com/teamtomo/libtilt). The project and a brief tutorial can be freely downloaded from github (https://github.com/Angeevillor/AI_HEAD).

## Acknowledgments

This work was supported in part by the National Key Research and Development Program of China (2021YFA1301500 to F. S. and 2021YFF0704300 to Y. Z.), the National Natural Science Foundation of China (32525005 and 92254306 to F. S., 32371248 to Y. Z.), and the Major Talent Project of Guangzhou National Laboratory (GZNL2025C01032 to D. Y. Z.). We are very grateful to Professor Edward H. Egelman from the University of Virginia in the United States for his full support and rigorous discussions in this work, and for providing multiple sets of laboratory data for testing.

