## Supplementary material for "AI-HEAD: Auto-indexing for Helical assembles’ diffraction and fast dynamic structure determination": All tables, videos and descriptions about the main

### Supplementary data

**Helical parameters for simulation results by auto-indexing**

| [n1,n2] | rise | P | twist | N | k_value | [Z1, Z2] | [x1_coff] | [x2_coff] |
| --- | --- | --- | --- | --- | --- | --- | --- | --- |
| [-1, 1] | 29.26 | 34.13 | 308.57 | 1 | [1, 0] | ['0.02930', '0.00488'] | [-1.0, 1.0, 0] | [1.0, -1.0, 1] |
| [-2, 1] | 25.6 | 204.8 | 45 | 1 | [0, 1] | ['0.02930', '0.00488'] | [-2.0, -2.0, 1] | [1.0, 1.0, 0] |
| [-1, 1] | 29.26 | 204.8 | 51.43 | 1 | [0, 1] | ['0.02930', '0.00488'] | [-1.0, -1.0, 1] | [1.0, 1.0, 0] |

**Table.S1** The solution set of helical parameters for indexing in Extend Figure. 2D (a.c). The correct solution for this set of indexing is in the main chart. The correct indexing is marked in red; Solutions in the additional chart are marked in green.

| [n1,n2] | rise | P | twist | N | k_value | [Z1, Z2] | [x1_coff] | [x2_coff] |
| --- | --- | --- | --- | --- | --- | --- | --- | --- |
| [-1, 1] | 25.6 | 204.8 | 45 | 1 | [0, 1] | ['0.03418', '0.00488'] | [-1.0, -1.0, 1] | [1.0,1.0,0] |
| [-1, 1] | 25.6 | 29.26 | 315 | 1 | [1, 0] | ['0.03418', '0.00488'] | [-1.0, 1.0, 0] | [1.0 -1.0,1] |

**Table.S2** The solution set of helical parameters for indexing in Extend Figure.2D(b). The correct solution for this set of indexing is in the additional chart The correct indexing is marked in red in the solution set, which is also in the additional chart.

**Helical parameters for EMDB data by auto-indexing**

| [n1, n2] | rise | P | twist | N | k_value | [Z1, Z2] | [x1_coff] | [x2_coff] |
| --- | --- | --- | --- | --- | --- | --- | --- | --- |
| [-1, 1] | 11.37 | 17.64 | 231.96 | 1 | [1, 0] | ['0.05669', '0.03129'] | [-1.0, 1.0, 0] | [1.0, -1.0, 1] |
| [-1, 2] | 6.91 | 17.64 | 141.07 | 1 | [1, 0] | ['0.05669', '0.03129'] | [-1.0, 1.0, 0] | [2.0, -2.0, 1] |
| [-1, 3] | 4.97 | 17.64 | 101.35 | 1 | [1, 0] | ['0.05669', '0.03129'] | [-1.0, 1.0, 0] | [3.0, -3.0, 1] |
| [-1, 4] | 3.88 | 17.64 | 79.09 | 1 | [1, 0] | ['0.05669', '0.03129'] | [-1.0, 1.0, 0] | [4.0, -4.0, 1] |
| [-2, 1] | 8.38 | 31.96 | 94.45 | 1 | [0, 1] | ['0.05669', '0.03129'] | [-2.0, -2.0, 1] | [1.0, 1.0, 0] |
| [-2, 2] | 11.37 | 35.28 | 115.98 | 2 | [1, 0] | ['0.05669', '0.03129'] | [-1.0, 1.0, 0] | [1.0, -1.0, 1] |
| [-2, 3] | 4.3 | 11.37 | 136.14 | 1 | [1, 1] | ['0.05669', '0.03129'] | [-2.0, -2.0, 1] | [3.0, 3.0, -1] |
| [-2, 4] | 6.91 | 35.28 | 70.53 | 2 | [1, 0] | ['0.05669', '0.03129'] | [-1.0, 1.0, 0] | [2.0, -2.0, 1] |
| [-1, 1] | 11.37 | 31.96 | 128.04 | 1 | [0, 1] | ['0.05669', '0.03129'] | [-1.0, -1.0, 1] | [1.0, 1.0, 0] |
| [-2, 2] | 11.37 | 63.91 | 64.02 | 2 | [0, 1] | ['0.05669', '0.03129'] | [-1.0, -1.0, 1] | [1.0, 1.0, 0] |

**Table.S3** The solution set of helical parameters for indexing of MAVS CARD in Extended Figure.3A. The correct solution for this set of indexing is in the main chart The correct indexing is marked in red in the solution set. Solutions in the additional chart are marked in green.

| [n1, n2] | rise | P | twist | N | k_value | [Z1, Z2] | [x1_coff] | [x2_coff] |
| --- | --- | --- | --- | --- | --- | --- | --- | --- |
| [-1, 1] | 17 | 67.56 | 90.6 | 1 | [1, 0] | ['0.01480', '0.04402'] | [-1.0, 1.0, 0] | [1.0, -1.0, 1] |
| [-2, 1] | 9.72 | 22.72 | 154.09 | 1 | [0, 1] | ['0.01480', '0.04402'] | [-2.0, -2.0, 1] | [1.0, 1.0, 0] |
| [-3, 1] | 6.81 | 22.72 | 107.9 | 1 | [0, 1] | ['0.01480', '0.04402'] | [-3.0, -3.0, 1] | [1.0, 1.0, 0] |
| [-4, 1] | 5.24 | 22.72 | 83.02 | 1 | [0, 1] | ['0.01480', '0.04402'] | [-4.0, -4.0, 1] | [1.0, 1.0, 0] |
| [-5, 1] | 4.26 | 22.72 | 67.46 | 1 | [0, 1] | ['0.01480', '0.04402'] | [-5.0, -5.0, 1] | [1.0, 1.0, 0] |
| [-6, 1] | 3.59 | 22.72 | 56.82 | 1 | [0, 1] | ['0.01480', '0.04402'] | [-6.0, -6.0, 1] | [1.0, 1.0, 0] |
| [-7, 1] | 3.1 | 22.72 | 49.07 | 1 | [0, 1] | ['0.01480', '0.04402'] | [-7.0, -7.0, 1] | [1.0, 1.0, 0] |
| [-8, 1] | 2.73 | 22.72 | 43.18 | 1 | [0, 1] | ['0.01480', '0.04402'] | [-8.0, -8.0, 1] | [1.0, 1.0, 0] |
| [-9, 1] | 2.43 | 22.72 | 38.56 | 1 | [0, 1] | ['0.01480', '0.04402'] | [-9.0, -9.0, 1] | [1.0, 1.0, 0] |
| [-10, 1] | 2.2 | 22.72 | 34.83 | 1 | [0, 1] | ['0.01480', '0.04402'] | [-10.0, -10.0, 1] | [1.0, 1.0, 0] |
| [-11, 1] | 2 | 22.72 | 31.76 | 1 | [0, 1] | ['0.01480', '0.04402'] | [-11.0, -11.0, 1] | [1.0, 1.0, 0] |
| [-12, 1] | 1.84 | 22.72 | 29.18 | 1 | [0, 1] | ['0.01480', '0.04402'] | [-12.0, -12.0, 1] | [1.0, 1.0, 0] |
| [-13, 1] | 1.7 | 22.72 | 26.99 | 1 | [0, 1] | ['0.01480', '0.04402'] | [-13.0, -13.0, 1] | [1.0, 1.0, 0] |
| [-14, 1] | 1.58 | 22.72 | 25.11 | 1 | [0, 1] | ['0.01480', '0.04402'] | [-14.0, -14.0, 1] | [1.0, 1.0, 0] |
| [-15, 1] | 1.48 | 22.72 | 23.47 | 1 | [0, 1] | ['0.01480', '0.04402'] | [-15.0, -15.0, 1] | [1.0, 1.0, 0] |
| [-16, 1] | 1.39 | 22.72 | 22.04 | 1 | [0, 1] | ['0.01480', '0.04402'] | [-16.0, -16.0, 1] | [1.0, 1.0, 0] |
| [-17, 1] | 1.31 | 22.72 | 20.77 | 1 | [0, 1] | ['0.01480', '0.04402'] | [-17.0, -17.0, 1] | [1.0, 1.0, 0] |
| [-18, 1] | 1.24 | 22.72 | 19.63 | 1 | [0, 1] | ['0.01480', '0.04402'] | [-18.0, -18.0, 1] | [1.0, 1.0, 0] |
| [-1, 1] | 17 | 22.72 | 269.4 | 1 | [0, 1] | ['0.01480', '0.04402'] | [-1.0, -1.0, 1] | [1.0, 1.0, 0] |

**Table.S4** The solution set of helical parameters for indexing of TMV in Extended Figure.3B. The correct solution for this set of indexing is in the main chart. The correct indexing is marked in red in the solution set. Solutions in the additional chart are marked in green.

| [n1, n2] | rise | P | twist | N | k_value | [Z1, Z2] | [x1_coff] | [x2_coff] |
| --- | --- | --- | --- | --- | --- | --- | --- | --- |
| [-1, 1] | 17.13 | 51.12 | 120.63 | 1 | [1, 0] | ['0.01956', '0.03882'] | [-1.0, 1.0, 0] | [1.0, -1.0, 1] |
| [-1, 2] | 12.83 | 51.12 | 90.35 | 1 | [1, 0] | ['0.01956', '0.03882'] | [-1.0, 1.0, 0] | [2.0, -2.0, 1] |
| [-1, 3] | 10.26 | 51.12 | 72.23 | 1 | [1, 0] | ['0.01956', '0.03882'] | [-1.0, 1.0, 0] | [3.0, -3.0, 1] |
| [-1, 4] | 8.54 | 51.12 | 60.16 | 1 | [1, 0] | ['0.01956', '0.03882'] | [-1.0, 1.0, 0] | [4.0, -4.0, 1] |
| [-2, 1] | 10.29 | 25.76 | 143.77 | 1 | [0, 1] | ['0.01956', '0.03882'] | [-2.0, -2.0, 1] | [1.0, 1.0, 0] |
| [-2, 2] | 17.13 | 102.24 | 60.31 | 2 | [1, 0] | ['0.01956', '0.03882'] | [-1.0, 1.0, 0] | [1.0, -1.0, 1] |
| [-2, 3] | 7.34 | 17.13 | 154.17 | 1 | [1, 1] | ['0.01956', '0.03882'] | [-2.0, -2.0, 1] | [3.0, 3.0, -1] |
| [-2, 4] | 12.83 | 102.24 | 45.18 | 2 | [1, 0] | ['0.01956', '0.03882'] | [-1.0, 1.0, 0] | [2.0, -2.0, 1] |
| [-3, 1] | 7.35 | 25.76 | 102.74 | 1 | [0, 1] | ['0.01956', '0.03882'] | [-3.0, -3.0, 1] | [1.0, 1.0, 0] |
| [-3, 2] | 6.43 | 17.13 | 135.09 | 1 | [1, 1] | ['0.01956', '0.03882'] | [-3.0, 3.0, -1] | [2.0, -2.0, 1] |
| [-3, 3] | 17.13 | 153.36 | 40.21 | 3 | [1, 0] | ['0.01956', '0.03882'] | [-1.0, 1.0, 0] | [1.0, -1.0, 1] |
| [-3, 4] | 5.14 | 17.13 | 107.94 | 1 | [1, 1] | ['0.01956', '0.03882'] | [-3.0, -3.0, 1] | [4.0, 4.0, -1] |
| [-4, 1] | 5.72 | 25.76 | 79.93 | 1 | [0, 1] | ['0.01956', '0.03882'] | [-4.0, -4.0, 1] | [1.0, 1.0, 0] |
| [-4, 2] | 10.29 | 51.52 | 71.89 | 2 | [0, 1] | ['0.01956', '0.03882'] | [-2.0, -2.0, 1] | [1.0, 1.0, 0] |
| [-4, 3] | 4.67 | 17.13 | 98.23 | 1 | [1, 1] | ['0.01956', '0.03882'] | [-4.0, 4.0, -1] | [3.0, -3.0, 1] |
| [-4, 4] | 17.13 | 204.48 | 30.16 | 4 | [1, 0] | ['0.01956', '0.03882'] | [-1.0, 1.0, 0] | [1.0, -1.0, 1] |
| [-1, 1] | 17.13 | 25.76 | 239.37 | 1 | [0, 1] | ['0.01956', '0.03882'] | [-1.0, -1.0, 1] | [1.0, 1.0, 0] |
| [-2, 2] | 17.13 | 51.52 | 119.69 | 2 | [0, 1] | ['0.01956', '0.03882'] | [-1.0, -1.0, 1] | [1.0, 1.0, 0] |
| [-3, 3] | 17.13 | 77.28 | 79.79 | 3 | [0, 1] | ['0.01956', '0.03882'] | [-1.0, -1.0, 1] | [1.0, 1.0, 0] |
| [-4, 4] | 17.13 | 103.05 | 59.84 | 4 | [0, 1] | ['0.01956', '0.03882'] | [-1.0, -1.0, 1] | [1.0, 1.0, 0] |

**Table.S5** The solution set of helical parameters for indexing of SNX1 complex in Extended Figure.3C. The correct solution for this set of indexing is in the main chart. The correct indexing is marked in red in the solution set. Solutions in the additional chart are marked in green.

| [n1,n2] | rise | P | twist | N | k_value | [Z1,Z2] | [x1_coff] | [x2_coff] |
| --- | --- | --- | --- | --- | --- | --- | --- | --- |
| [-1, 1] | 24.44 | 41.35 | 212.73 | 1 | [1, 0] | ['0.02418', '0.01674'] | [-1.0, 1.0, 0] | [1.0, -1.0, 1] |
| [-1, 2] | 15.36 | 41.35 | 133.71 | 1 | [1, 0] | ['0.02418', '0.01674'] | [-1.0, 1.0, 0] | [2.0, -2.0, 1] |
| [-1, 3] | 11.2 | 41.35 | 97.5 | 1 | [1, 0] | ['0.02418', '0.01674'] | [-1.0, 1.0, 0] | [3.0, -3.0, 1] |
| [-1, 4] | 8.81 | 41.35 | 76.72 | 1 | [1, 0] | ['0.02418', '0.01674'] | [-1.0, 1.0, 0] | [4.0, -4.0, 1] |
| [-1, 5] | 7.26 | 41.35 | 63.24 | 1 | [1, 0] | ['0.02418', '0.01674'] | [-1.0, 1.0, 0] | [5.0, -5.0, 1] |
| [-1, 6] | 6.18 | 41.35 | 53.79 | 1 | [1, 0] | ['0.02418', '0.01674'] | [-1.0, 1.0, 0] | [6.0, -6.0, 1] |
| [-1, 7] | 5.38 | 41.35 | 46.8 | 1 | [1, 0] | ['0.02418', '0.01674'] | [-1.0, 1.0, 0] | [7.0, -7.0, 1] |
| [-1, 8] | 4.76 | 41.35 | 41.42 | 1 | [1, 0] | ['0.02418', '0.01674'] | [-1.0, 1.0, 0] | [8.0, -8.0, 1] |
| [-1, 9] | 4.27 | 41.35 | 37.14 | 1 | [1, 0] | ['0.02418', '0.01674'] | [-1.0, 1.0, 0] | [9.0, -9.0, 1] |
| [-1, 10] | 3.87 | 41.35 | 33.67 | 1 | [1, 0] | ['0.02418', '0.01674'] | [-1.0, 1.0, 0] | [10.0, -10.0, 1] |
| [-1, 11] | 3.54 | 41.35 | 30.79 | 1 | [1, 0] | ['0.02418', '0.01674'] | [-1.0, 1.0, 0] | [11.0, -11.0, 1] |
| [-1, 12] | 3.26 | 41.35 | 28.36 | 1 | [1, 0] | ['0.02418', '0.01674'] | [-1.0, 1.0, 0] | [12.0, -12.0, 1] |
| [-1, 13] | 3.02 | 41.35 | 26.29 | 1 | [1, 0] | ['0.02418', '0.01674'] | [-1.0, 1.0, 0] | [13.0, -13.0, 1] |
| [-1, 14] | 2.81 | 41.35 | 24.5 | 1 | [1, 0] | ['0.02418', '0.01674'] | [-1.0, 1.0, 0] | [14.0, -14.0, 1] |
| [-2, 1] | 17.34 | 59.73 | 104.52 | 1 | [0, 1] | ['0.02418', '0.01674'] | [-2.0, -2.0, 1] | [1.0, 1.0, 0] |
| [-2, 2] | 24.44 | 82.71 | 106.36 | 2 | [1, 0] | ['0.02418', '0.01674'] | [-1.0, 1.0, 0] | [1.0, -1.0, 1] |
| [-2, 3] | 9.43 | 24.44 | 138.95 | 1 | [1, 1] | ['0.02418', '0.01674'] | [-2.0, -2.0, 1] | [3.0, 3.0, -1] |
| [-2, 4] | 15.36 | 82.71 | 66.86 | 2 | [1, 0] | ['0.02418', '0.01674'] | [-1.0, 1.0, 0] | [2.0, -2.0, 1] |
| [-2, 5] | 6.48 | 15.36 | 151.81 | 1 | [2, 1] | ['0.02418', '0.01674'] | [-2.0, -2.0, 1] | [5.0, 5.0, -2] |
| [-2, 6] | 11.2 | 82.71 | 48.75 | 2 | [1, 0] | ['0.02418', '0.01674'] | [-1.0, 1.0, 0] | [3.0, -3.0, 1] |
| [-2, 7] | 4.93 | 11.2 | 158.53 | 1 | [3, 1] | ['0.02418', '0.01674'] | [-2.0, -2.0, 1] | [7.0, 7.0, -3] |
| [-2, 8] | 8.81 | 82.71 | 38.36 | 2 | [1, 0] | ['0.02418', '0.01674'] | [-1.0, 1.0, 0] | [4.0, -4.0, 1] |
| [-2, 9] | 3.98 | 8.81 | 162.67 | 1 | [4, 1] | ['0.02418', '0.01674'] | [-2.0, -2.0, 1] | [9.0, 9.0, -4] |
| [-2, 10] | 7.26 | 82.71 | 31.62 | 2 | [1, 0] | ['0.02418', '0.01674'] | [-1.0, 1.0, 0] | [5.0, -5.0, 1] |
| [-2, 11] | 3.34 | 7.26 | 165.47 | 1 | [5, 1] | ['0.02418', '0.01674'] | [-2.0, -2.0, 1] | [11.0, 11.0, -5] |
| [-2, 12] | 6.18 | 82.71 | 26.9 | 2 | [1, 0] | ['0.02418', '0.01674'] | [-1.0, 1.0, 0] | [6.0, -6.0, 1] |
| [-2, 13] | 2.87 | 6.18 | 167.49 | 1 | [6, 1] | ['0.02418', '0.01674'] | [-2.0, -2.0, 1] | [13.0, 13.0, -6] |
| [-2, 14] | 5.38 | 82.71 | 23.4 | 2 | [1, 0] | ['0.02418', '0.01674'] | [-1.0, 1.0, 0] | [7.0, -7.0, 1] |
| [-3, 1] | 13.44 | 59.73 | 81 | 1 | [0, 1] | ['0.02418', '0.01674'] | [-3.0, -3.0, 1] | [1.0, 1.0, 0] |
| [-3, 2] | 10.14 | 24.44 | 149.43 | 1 | [1, 1] | ['0.02418', '0.01674'] | [-3.0, 3.0, -1] | [2.0, -2.0, 1] |
| [-3, 3] | 24.44 | 124.06 | 70.91 | 3 | [1, 0] | ['0.02418', '0.01674'] | [-1.0, 1.0, 0] | [1.0, -1.0, 1] |
| [-3, 4] | 6.81 | 24.44 | 100.25 | 1 | [1, 1] | ['0.02418', '0.01674'] | [-3.0, -3.0, 1] | [4.0, 4.0, -1] |
| [-3, 5] | 5.84 | 15.36 | 136.96 | 1 | [-2, -1] | ['0.02418', '0.01674'] | [-3.0, 3.0, -1] | [5.0, -5.0, 2] |
| [-3, 6] | 15.36 | 124.06 | 44.57 | 3 | [1, 0] | ['0.02418', '0.01674'] | [-1.0, 1.0, 0] | [2.0, -2.0, 1] |
| [-3, 7] | 4.56 | 15.36 | 106.78 | 1 | [2, 1] | ['0.02418', '0.01674'] | [-3.0, -3.0, 1] | [7.0, 7.0, -2] |
| [-3, 8] | 4.1 | 11.2 | 131.91 | 1 | [-3, -1] | ['0.02418', '0.01674'] | [-3.0, 3.0, -1] | [8.0, -8.0, 3] |
| [-3, 9] | 11.2 | 124.06 | 32.5 | 3 | [1, 0] | ['0.02418', '0.01674'] | [-1.0, 1.0, 0] | [3.0, -3.0, 1] |
| [-3, 10] | 3.42 | 11.2 | 110.06 | 1 | [3, 1] | ['0.02418', '0.01674'] | [-3.0, -3.0, 1] | [10.0, 10.0, -3] |
| [-3, 11] | 3.16 | 8.81 | 129.18 | 1 | [-4, -1] | ['0.02418', '0.01674'] | [-3.0, 3.0, -1] | [11.0, -11.0, 4] |
| [-3, 12] | 8.81 | 124.06 | 25.57 | 3 | [1, 0] | ['0.02418', '0.01674'] | [-1.0, 1.0, 0] | [4.0, -4.0, 1] |
| [-3, 13] | 2.74 | 8.81 | 112.04 | 1 | [4, 1] | ['0.02418', '0.01674'] | [-3.0, -3.0, 1] | [13.0, 13.0, -4] |
| [-3, 14] | 2.57 | 7.26 | 127.46 | 1 | [-5, -1] | ['0.02418', '0.01674'] | [-3.0, 3.0, -1] | [14.0, -14.0, 5] |
| [-4, 1] | 10.97 | 59.73 | 66.12 | 1 | [0, 1] | ['0.02418', '0.01674'] | [-4.0, -4.0, 1] | [1.0, 1.0, 0] |
| [-4, 2] | 17.34 | 119.47 | 52.26 | 2 | [0, 1] | ['0.02418', '0.01674'] | [-2.0, -2.0, 1] | [1.0, 1.0, 0] |
| [-4, 3] | 7.17 | 24.44 | 105.6 | 1 | [1, 1] | ['0.02418', '0.01674'] | [-4.0, 4.0, -1] | [3.0, -3.0, 1] |
| [-4, 4] | 24.44 | 165.42 | 53.18 | 4 | [1, 0] | ['0.02418', '0.01674'] | [-1.0, 1.0, 0] | [1.0, -1.0, 1] |
| [-4, 5] | 5.32 | 24.44 | 78.42 | 1 | [1, 1] | ['0.02418', '0.01674'] | [-4.0, -4.0, 1] | [5.0, 5.0, -1] |
| [-4, 6] | 9.43 | 48.87 | 69.47 | 2 | [1, 1] | ['0.02418', '0.01674'] | [-2.0, -2.0, 1] | [3.0, 3.0, -1] |
| [-4, 7] | 4.23 | 15.36 | 99.21 | 1 | [-2, -1] | ['0.02418', '0.01674'] | [-4.0, 4.0, -1] | [7.0, -7.0, 2] |
| [-4, 8] | 15.36 | 165.42 | 33.43 | 4 | [1, 0] | ['0.02418', '0.01674'] | [-1.0, 1.0, 0] | [2.0, -2.0, 1] |
| [-4, 9] | 3.51 | 15.36 | 82.35 | 1 | [2, 1] | ['0.02418', '0.01674'] | [-4.0, -4.0, 1] | [9.0, 9.0, -2] |
| [-4, 10] | 6.48 | 30.72 | 75.9 | 2 | [2, 1] | ['0.02418', '0.01674'] | [-2.0, -2.0, 1] | [5.0, 5.0, -2] |
| [-4, 11] | 3 | 11.2 | 96.54 | 1 | [-3, -1] | ['0.02418', '0.01674'] | [-4.0, 4.0, -1] | [11.0, -11.0, 3] |
| [-4, 12] | 11.2 | 165.42 | 24.38 | 4 | [1, 0] | ['0.02418', '0.01674'] | [-1.0, 1.0, 0] | [3.0, -3.0, 1] |
| [-4, 13] | 2.62 | 11.2 | 84.29 | 1 | [3, 1] | ['0.02418', '0.01674'] | [-4.0, -4.0, 1] | [13.0, 13.0, -3] |
| [-4, 14] | 4.93 | 22.4 | 79.27 | 2 | [3, 1] | ['0.02418', '0.01674'] | [-2.0, -2.0, 1] | [7.0, 7.0, -3] |
| [-1, 1] | 24.44 | 59.73 | 147.27 | 1 | [0, 1] | ['0.02418', '0.01674'] | [-1.0, -1.0, 1] | [1.0, 1.0, 0] |
| [-2, 2] | 24.44 | 119.47 | 73.64 | 2 | [0, 1] | ['0.02418', '0.01674'] | [-1.0, -1.0, 1] | [1.0, 1.0, 0] |
| [-3, 3] | 24.44 | 179.2 | 49.09 | 3 | [0, 1] | ['0.02418', '0.01674'] | [-1.0, -1.0, 1] | [1.0, 1.0, 0] |
| [-4, 4] | 24.44 | 238.93 | 36.82 | 4 | [0, 1] | ['0.02418', '0.01674'] | [-1.0, -1.0, 1] | [1.0, 1.0, 0] |

**Table.S6** The solution set of helical parameters for indexing of SIRV2 in Extended Figure.3D. The correct solution for this set of indexing is in the main chart. The correct indexing is marked in red in the solution set. Solutions in the additional chart are marked in green.

**Visualization of parameter sorting by diffraction pattern using AI-HEAD**

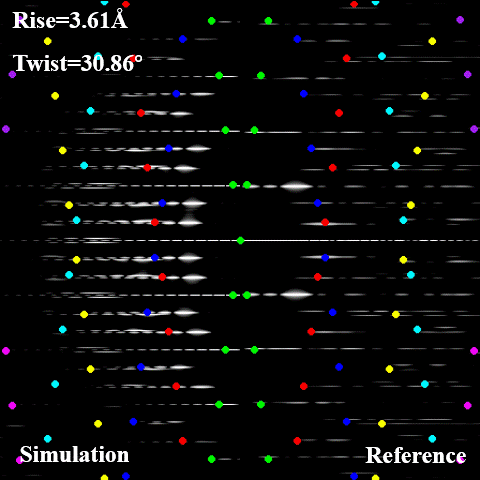

**Video. S1** The simulated lattices in Diffraction-sorting of SIRV2 which are similar to the result of "Auto-indexing". The correct parameters are highlighted in red.

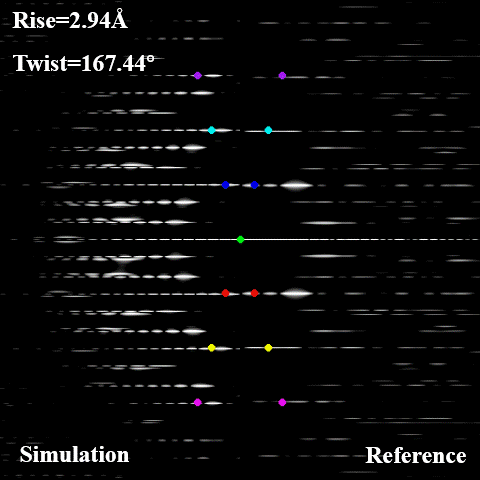

**Video. S2** The simulated lattices in Diffraction-sorting of SIRV2 which are similar to a part of the result of "Auto-indexing".

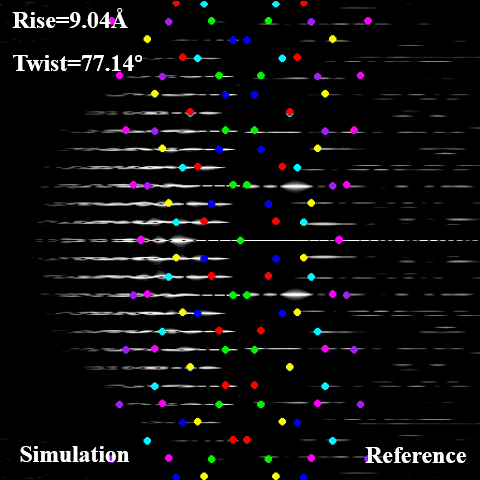

**Video. S3** The simulated lattices in Diffraction-sorting of SIRV2 which are not similar to the result of "Auto-indexing". These parameters can be removed during the calculation.

**Design and Specifications of Each Module**

##### Image_information

This module is designed to determine the diffraction center of the diffraction pattern and to obtain the location of the diffraction patches. This step is prepared for the “Global searching” in “Solve_FFT” module. The user could load the screenshot of the diffraction pattern as input. Since the center of the captured diffraction pattern is not necessarily the exact center of the image, the program will first display the diffraction pattern, and users can specify the location of the diffraction center by clicking through the interactive interface. The program automatically writes the coordinates of the diffraction center to the center's log file. As only the major diffraction patches and black background are left in the diffraction pattern image, there is a large contrast between the diffraction signal and the background, and all the regions of the diffraction signal can be found by edge detection. AI-HEAD use Canny algorithm (Canny, 1986) to detect the diffraction signal first, and then OSTU algorithm is used to calculate the optimal segmentation threshold and realize automatic segmentation (N., 1979; A. and S. et al., 2018; D. and H. et al., 2018; Latychevskaia and Fink, 2018; N. and C., 2018; P. and K., 2018). The coordinates of all detected diffraction patches are automatically written to the region log file, and all of the selected region is marked with a rectangle in pattern and exported. Sometimes the algorithm may not be able to find all the diffraction patches, or for the average diffraction pattern, some of the diffraction patches may fuse together and not be well separated. AI-HEAD provides a manual correction step: after edge detection, AI-HEAD provides an interactive interface again, whereby the user can manually supplement and split the existing diffraction pattern according to the result of edge detection to ensure that the coordinate information of all diffraction patches in the pattern is obtained. All of the results are automatically saved under the “Image_information” path.

##### Solve_FFT

This module is designed to determine the position of optimal basic vectors by using “Global searching” strategy. The user needs to load the diffraction pattern along with log file got in “Image_information” module to get the coordinate of diffraction center and diffraction region, and input the number of optimal solutions that need to be output. Then, AI-HEAD will extract all of the center coordinate of diffraction patches, which transformed the diffraction patches into diffraction points. As mentioned earlier, Fourier space is divided into four quadrants by the equator and the meridian. The program will automatically construct a basic vector for all the diffraction points in the first quadrant one by one as a set of basic vectors whose n are positive, and perform the same operation for the diffraction points in the second quadrant to obtain a set of basic vectors whose n are negative. Next, the program will take one vector from each of the two sets to form a pair of vectors one by one, and start from the center of diffraction pattern to produce lattice points and symmetric lattice points. In this way, the complete “solution set” of the basic vectors is obtained. The program then sorts the “solution set” according to the conditions of the ideal indexing: First, the program determines whether there are diffraction patches near the meridian. If there is a diffraction patch, the region of the lattice produced by the basic vectors near the meridian can only fall on the diffraction plate, but not in other regions. If this condition is not met, it is not an ideal solution. If there is no diffraction patch near the meridian, the lattice generated by the basic vectors cannot fall in the region near the meridian, otherwise it is not an ideal solution. In the case of confirming that the solution is ideal, the program automatically delimits the coordinate region of the diffraction layer line according to the coordinate region of the diffraction patch which is used to compares the number of diffraction layer lines occupied by the resulting lattice. Finally, for solutions that occupy the same number of diffraction layer lines, the program again orders them again according to the number of diffraction patches covered by the lattice. This completes the sorting of the entire solution set, and then the program outputs the first n solutions according to the number specified by the user and automatically outputs them to the program directory. At the same time, the program will automatically output the sorted data into a table, which records the number of diffraction layer lines of the output solution, the number of points on each diffraction layer and the number of diffraction patches covered. All of the results are automatically saved under the "Solve_FFT " path.

##### Solve_FFT_Refine

This module is designed to determine the position of optimal basic vectors by using “Local refinement” strategy. The user needs to load the diffraction pattern along with log file got in Image_information module to get the coordinate of diffraction center and diffraction region. As the “Global searching” strategy only takes the center point of each diffraction patch as the diffraction point, there may be a large error when the diffraction patch is long. Therefore, AI-HEAD provides the FAST algorithm(E. and R. et al., 2010), which is able to detect more feature points on a diffraction spot. The user can adjust different detection modes and detection thresholds to obtain the appropriate number of feature points. Then the user can select a certain sampling area in the first and second quadrants respectively on the interactive interface (it is not recommended to delimit a large area, otherwise it will lead to a slow operation speed), and the program will perform the same work as the global search. Then the solution set is sorted and output according to the number of covered diffraction points. The main effect of this step is that the location of the diffraction point may not be in the center of the diffraction patch in the optimal solution for indexing, and more precise location can be found by “Local refinement”. All of the results are automatically saved under the "Solve_FFT_Refine" path.

##### Basic_vector_cooordinate:

This module is designed to get the diffraction pattern and determine the coordinate of optimal basic vectors in Fourier space based on Fourier-Bessel analysis. The user needs to load the original data file (in mrc format), which could be the tube or the power spectrum (2D average power spectrum is recommended). AI-HEAD provide an interactive interface to display the original data. The interactive interface provides three sliders, which can adjust the magnification, contrast and brightness of image respectively. The user could get the diffraction pattern by adjusting the contrast and brightness to preserve the main diffraction signal and remove the excess noise. If the user uses this module to obtain diffraction pattern, the adjusted pattern screenshot can be load in Image_information module and conduct the indexing process. If the user has obtained the position of the optimal basic vector on the diffraction pattern, then only need to find the diffraction point on the interactive interface, by double-clicking the diffraction point, the program will automatically convert the pixel coordinates of the base vector to n-z coordinates and write in the logfile.

According to the Fourier-Bessel principle, the diffraction intensity on the n-order diffraction layer is proportional to the n-order Bessel function(Hawkes and Valdré, 1991; Ward and Moody et al., 2003; Diaz and Rice et al., 2010; Egelman, 2010; Zhang, 2013). The horizontal coordinate of the maximum point of the Bessel function and the order of the Bessel function satisfies the following statistical relationship(Zhang, 2013):

$$2\pi Rr=1.03\left| n \right|+1$$

Where R represents the distance from the diffraction point to the meridian, r represents the helical radius (outer diameter), and n represents the Bessel order. Through this statistical relationship, the Bessel order corresponding to the basic vectors can be roughly calculated. All of the results are automatically saved under the “Basic_vector_cooordinate” path.

##### Helical_parameter:

This module is designed to calculate the all of the possible helical parameters as a chart based on Fourier-Bessel analysis. The user needs to load the coordinate of two basic vectors got in Basic_vector_cooordinate module. According to Fourier-Bessel analysis, the greatest common divisor of the Bessel order of two basis vectors is the minimum starting number of the helix, which is the C_n_ symmetry of the helix(Zhang, 2013). If the coordinates of two basic vectors are (n_1_ (negative), z_1_) and (n_2_ (positive), z_2_), and N is denoted as the greatest common factor of n_1_ and n2, then the helix has C_n_ symmetric. Since the continuous helical is equivalent to the convolution continuation of the helical density in units of period. So, in Fourier space, helical diffraction is sampled with the reciprocal of the periodic distance. In other words, the z coordinate of the first-order diffraction layer line is the pitch (P) of the helix, only the point located on the first-order diffraction layer line needs to be represented by the basic vectors, and the pitch can be obtained according to the coordinate information.

$P=\frac{N}{{|k}_{1}z_{1}+k_{2}z_{2}|}$ ($\left| k_{1}n_{1}+k_{2}n_{2} \right|=N$)

Since the helical assembly is not continuous, it is equivalent to sampling the continuous helix in real space, and the sampling distance in Fourier space is equal to rise. Therefore, the basic vector is used to represent the first diffraction point above the diffraction center along the meridian direction. and the rise ($\Delta z$) can be obtained according to the coordinate information.

$\Delta z=\frac{N}{{|n}_{2}z_{1}-n_{1}z_{2}|}$

And then, the twist ($\Delta\varphi$) of the helix could be calculated by the pitch and rise.

$$\Delta\varphi=\frac{{360}^{^{\circ}}\Delta z}{P}$$

Since the helical assembly has inner and outer diameter, the Bessel order of vector coordinates estimated by calculating the outer diameter is actually the maximum of the Bessel order of the vector. In other words, if the Bessel order of vector_1_ is n_1_ and the Bessel order of vector_2_ is n_2_, then there are n_1_*n_2_ combinations of basis vectors. And when the absolute values of n_1_ and n_2_ are the same, there are two different scenarios for k_1_ and k_2_. So, an additional n (min (|n_1_|, n_2_)) results are written in the “additional chart” (See in Table S1-2 in detail). Details on the interpretation of data in each column of the parameter list can be provided in supplemental data. All of the results are automatically saved under the “Helix_parameter” path.

##### Helix_simulation:

This module is designed to simulate the helix based the all of the possible helical parameters. The user needs to load the chart of helical parameters obtained from the “Helix_parameter” module and determine the pixel size, the length and radius of the simulated helix. The program will automatically read the helical parameters in the chart and generates a series of projection based on these parameters. Meanwhile, based on each set of parameters, the program could also calculate the diffraction information of the simulated projections in the reciprocal space and writes it to the log file for later diffraction sorting. All of the results are automatically saved in the “projection” folder under the “Helix_simulation” path.

##### Diffraction_simulation:

This module is designed to obtain diffraction patterns based on the helical projections in batch. There are “single” and “All” modes in this module. Since it is required to adjust the diffraction power spectrum before obtaining the diffraction pattern, the user can first adjust one of the projections in the “single” mode to obtain a more suitable diffraction pattern. Meanwhile, information such as the adjusted brightness, contrast, and the region of diffraction pattern can be exported to a “value” file. Next, the user can switch to the “All” mode, and the program will automatically diffract all the remaining projections according to the information adjusted in the “single” mode and output them in batch. The user could modify the data in the “value” file to achieve different adjustment methods until a more ideal diffraction pattern is obtained. All of the results are automatically saved under the “Diffraction_simulation” path.

##### Diffraction_sorting:

This module is designed to compare the actual diffraction pattern with the simulated diffraction pattern based on the helical parameters of simulation data. The user could use the real diffraction pattern as a reference, and then pair the generated simulation data with the original data to form a series of “converge images” in turn. All “converge images” will be saved in “converge” folder automatically under the " Diffraction_sorting " path. Next, the program will generate the simulation lattice according to the helical parameters of the simulated data. The ideal diffraction pattern consists of a series of equally spaced "diffraction layer lines", and the intensity on each layer obeys the law of a certain order Bessel function, and extends up and down along the meridian according to a certain distance (See Extended Figure.1). Therefore, AI-HEAD uses different colors to distinguish points from patterns on different diffraction centers: (Brown: center located on (0, 4/Δz), Purple: center located on (0, 3/Δz), Cyan: center located on (0, 2/Δz), Blue: center located on (0, 1/Δz), Green: center located on (0,0), Red: center located on (0, -1/Δz), Yellow: center located on (0, -2/Δz), Pink: center located on (0, -3/Δz) Chocolate: (0, -4/Δz). Finally, the user can filter out part of the solution set according to the comparison result, thus simplifying the subsequent filtering process. All of the results are automatically saved in the “compare” folder under the “Diffraction_sorting” path.

##### Extract_particles:

This module is designed to extract tubes or filaments from electron microscopy images and segment them into discrete particles. AI-HEAD is capable of reading coordinates picked by e2helixboxer.py, and performing batch extraction of tubes or filaments based on user-defined widths. Furthermore, the extracted tubes and filaments could be preliminarily categorized into distinct subclasses using subsequent diameter-based classification modules. All of the results are automatically saved under the “Extract_particles” path.

##### Diameter_classification:

This module is designed to quantitatively assess the diameters of filaments or tubes and to classify them into distinct subgroups based on their outer diameters, as defined by the user-specified number of classes. The diameter estimation procedure involves projecting each structure along the y-axis to generate a one-dimensional intensity profile. The outer and inner diameters are determined by measuring the distance between the two principal peaks in the projection curve, corresponding to the structural boundaries. These measurements are subsequently used to assign each filament or tube to a diameter-based subclass, facilitating downstream structural or functional analyses. All of the results are automatically saved under the “Diameter_classification” path.

##### Sorting_coordinates:

This module is designed to filter out the coordinates of tubes of a certain diameter class from all the coordinates of tubes and export them in the EMAN2 format. In this way, it is possible to extract and analyze tubes or filaments of a certain diameter separately, and obtain particles at a specific diameter. All of the results are automatically saved under the “Sorting_coordinates” path.

##### Average_power_spectra:

This module is designed to obtain the average power spectra of the entire class by averaging the power spectrum of a specific type of tubes or filaments. The filaments or tubes are first cut to a specific size, then padded to the specified size (in general 2048 or 4096). Following Fourier transformation, the resulting power spectra are averaged to yield a representative spectrum for the specific class. All of the results are automatically saved under the “Average_power_spectra” path.

##### Generate_initial_model

This module is designed to generate feature-free cylinders as the initial model by SPIDER (Egelman, 2007). All of the results are automatically saved under the “Generate_initial” path.

##### IHRSR

This module is designed to generate projects of IHRSR based on the helical parameters calculated before. By running the IHRSR project in turn, the only correct helical parameter among a series of candidate parameters can be obtained, as well as a better reference. All of the results are automatically saved under the “IHRSR” path.
